# Heavy and wet: evaluating the validity and implications of assumptions made when measuring growth efficiency using ^18^O water

**DOI:** 10.1101/601138

**Authors:** Grace Pold, Luiz A. Domeignoz-Horta, Kristen M. DeAngelis

## Abstract

How microbes allocate carbon to growth vs. respiration plays a central role in determining the ability of soil to retain carbon. This carbon use efficiency (CUE) is increasingly measured using the ^18^O-H_2_O method, in which heavy oxygen incorporated into DNA is used to estimate growth. Here we evaluated the validity of some of the assumptions of this method using a literature search, and then tested how violating them affected estimates of the growth component of carbon use efficiency in soil. We found that the ^18^O method is consistently sensitive to assumptions made about oxygen sources to DNA, but that the effect of other assumptions depends on the microbial community present. We provide an example for how the tools developed here may be used with observed CUE values, and demonstrate that the original conclusions drawn from the data remain robust in the face of methodological bias. Our results lay the foundation for a better understanding of the consequences to the ^18^O method underlying assumptions. Future studies can use the approach developed here to identify how different incubation conditions and/or treatments might bias its CUE estimates and how trustworthy their results are. Further wet-lab work dissecting the assumptions of the ^18^O method in soil will help justify the scenarios under which it is reasonable to trust its results.

## 2 Introduction

Carbon use efficiency - or the fraction of carbon taken up by a cell and retained in biomass - is a central determinant of soil organic matter longevity. Across a wide range of complexities, models of the carbon cycle have shown that the degree to which soil organic matter is lost in a warmer world is contingent upon carbon use efficiency [1, 2, 3, 4]. For many years, the study of CUE was limited to looking at one substrate type at a time, as a single heavy-labeled carbon source was added to the soil. Under this method, heavy carbon is partitioned by the cell into respiration and biomass, and carbon use efficiency can be calculated as the fraction of heavy carbon collected from biomass compared to the sum collected from biomass and CO_2_ respiration. However, this method is believed to overestimate “true” efficiency by measuring the uptake of simple labile compounds, and not their integration into biomass [5, 6]. ^13^C methods may also overestimate CUE if the target compound preferentially enters anabolic pathways while non-labeled substrates are used to generate ATP [7, 8]. Finally, ^13^C methods measure substrate use efficiency on a specific compound, and do not capture the repertoire of substrates microbes are faced with in natural environments such as soil.

Due to these known biases there has been a recent push towards using substrate-agnostic growth-based measures of biomass increment, such as ^18^O-H_2_O incorporation into DNA. This method provides more realistic and reproducible measures of CUE than the other dominant methods [5]. To complete this assay, ^18^O-H_2_O is added at 5-50% of the total soil moisture and the soil is incubated in a sealed container for 12-72 hours [5, 9]. At the end of the incubation, a gas sample is taken to measure the dissimilatory carbon losses, and the incubated soil is extracted for DNA. The amount of ^18^O incorporated into the DNA is then determined using Isotope-Ratio Mass Spectrometry (IRMS), and converted into new DNA produced assuming 31% of DNA is oxygen. This DNA “growth” is then converted into microbial biomass carbon produced using either a sample-specific [5, 10, 11] or a cross-sample average ratio between total DNA yield and chloroform fumigation-extractable microbial biomass carbon [12]. CUE can then be calculated as for the ^13^C and ^14^C labeled methods.

For the ^18^O-H_2_O method to provide an accurate estimate of CUE, a number of assumptions must be made. These include extracellular water being the sole source of oxygen in DNA; unbiased DNA and microbial biomass carbon extraction, and the actively growing community being representative of the total community (Figure 1). Here we explore the validity of these assumptions, the effects of violating them, and the subsequent consequences for the conclusions made. We focus primarily on how the sensitivity of conclusions changes as a function of the fungal:bacterial DNA ratio of soil, both because proxies for this value are often determined during routine soil analyses, and because relevant physiological differences between these two groups are relatively well studied.

**Figure 1:**
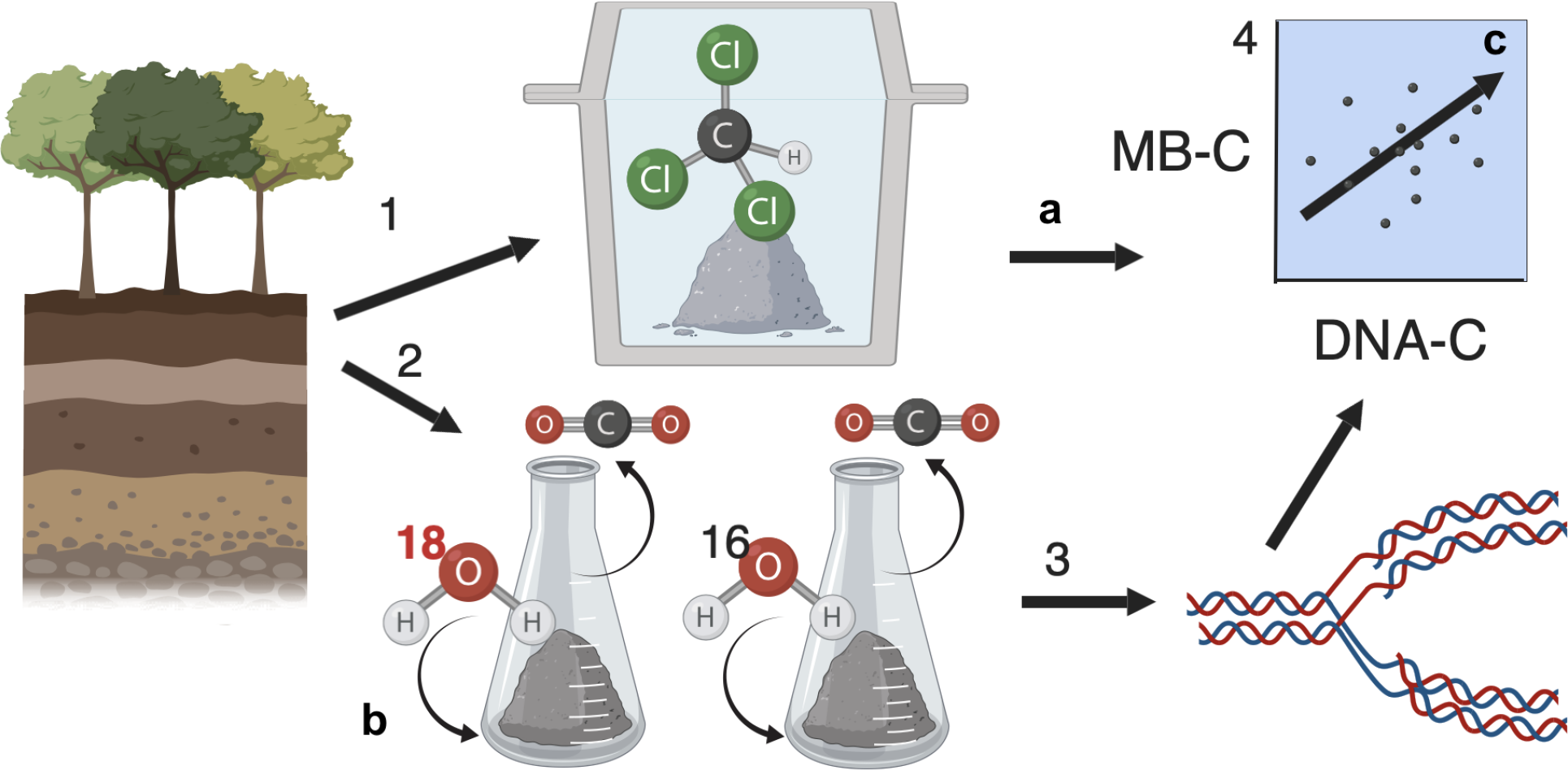
The ^18^O-H_2_O method of evaluating microbial growth in units of carbon (numbers), and the assumptions made (letters). 1. Soil collected from the environment is subject to chloroform fumigation extraction to determine total microbial biomass carbon. All taxa are assumed to have their biomass extracted with equal and complete efficiency (a). 2. A subfraction of the soil is incubated with ^18^O-H_2_O, which is assumed to be incorporated into new DNA (3) to comprise a fraction of the oxygens equal to its abundance as a fraction of total soil water (b). 4. The DNA is extracted and quantified, so that a relationship between the DNA and microbial biomass carbon content of the community can be established. It is assumed that this community-level MBC:DNA ratio is representative of the community which grew during the incubation with ^18^O-H_2_O, such that the new DNA growth can be converted to new microbial biomass carbon (c). Image made in BioRender©- biorender.com

## 3 Methods

We generated models to simulate the effects of inefficient DNA and MBC extraction, the active community not representing the total community, alternative oxygen sources to DNA, and differential growth rates between bacteria and fungi on measured MBC accumulation. All analyses were completed in R v3.4.0 [13], and results were plotted using ggplot2 [14]. Other packages used for the analysis included: plyr [15], Shiny [16], and ggpubr [17].

### 3.1 Model development

First, we explored the existing literature for reported values regarding each parameter corresponding to the underlying assumptions that could impact CUE estimates (Table 1). Second, we generated a Shiny app [16] to interactively explore the effect of violating the assumptions of microbial growth measurements over a range of fungal:bacterial ratios. It is available at: https://gracepold.shinyapps.io/18OSimulations/ until 25 hours/month server time have been used, and also as supplementary file S1. These simulations were run either assuming identical growth rates for bacteria and fungi (which were generated as a function of the fungal:bacterial DNA ratio), or that bacteria and fungi formed groups with distinct growth rates. Within each of these scenarios, we evaluated subsets where just DNA was extracted inefficiently, where MBC was extracted inefficiently, or where both were incompletely extracted. This app was additionally used for error checking the code used in subsequent steps, as predicted responses and test cases could be readily screened than when embedded in sensitivity analyses.

**Table 1:** Values used to parameterize simulations of microbial biomass carbon growth during ^18^O-H_2_O addition to soil. See File S2 for full details

| Definition | Parameter | value | minimum | maximum | Reference |
| --- | --- | --- | --- | --- | --- |
| Metabolically active fraction bacteria | activeFractionB | 0.1 | 0.01 | 1 | [23, 24, 25] |
| Metabolically active fraction of fungi | activeFractionF | 0.1 | 0.01 | 1 | NA |
| Fraction of DNA oxygen from extracellular water | DNAbias | 0.7 | 0.3 | 1 | [5, 26, 27, 28, 29, 30] |
| Bacterial DNA extraction efficiency | DNAexteffB | 0.5 | 0.06 | 0.5 | [31], personal obs |
| Fungal DNA extraction efficiency | DNAexteffF | 0.33 | 0.04 | 0.5 | [31], personal obs |
| Bacterial growth rate (daily multiplier) | GRbact | 0.167 | 0.003 | 3.4 | [32, 33, 34, 35, 36] |
| Fungal growth rate (daily multiplier) | GRfun | 0.0088 | 0.0001 | 0.8 | [32, 33, 37, 35], Eric Morrison, pers. comm. |
| MBC:DNA ratio of bacteria | MBCDNAratB | 13.22 | 3.6 | 37 | [38, 39, 40, 41, 42, 43, 44, 35] |
| MBC:DNA ratio of fungi | MBCDNAratF | 1070 | 90 | 3300 | [45, 39, 41, 46, 47] |
| Bacterial MBC extraction efficiency | MBCexteffB | 0.32 | 0.1 | 0.51 | [48, 49, 50, 51] |
| Fungal MBC extraction efficiency | MBCexteffF | 0.33 | 0.21 | 0.45 | [49, 50, 51, 52] |
| Bacterial 16S copies per genome | rrnpergenome | 2.25 | 1 | 15 | [53, 22] |
| Fungal ITS copies per genome | ITSpergenome | 82 | 14 | 1442 | [21] |

Next, we completed a sensitivity analysis by running simulations where a single parameter was changed to the minimum or maximum value observed in the literature (supplementary file S1 and 1), while keeping all the remaining parameters at a best-estimate value. Since “true” MBC differed between simulations, we divided the resultant “apparent” or “observed” microbial biomass carbon by the true microbial biomass carbon in order to standardize results. Sensitivity values were subsequently recorded as per Allison *et al*. [1, 6]:

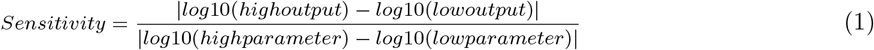

where *high output* is the ratio of the true CUE to the observed CUE under the high parameter value, and *low output* is the ratio of the observed CUE under the low parameter value. Simulation parameters and the underlying assumptions can be found in Table 1, and references are available as a part of our Shiny app.

### 3.2 Empirical validation

We used a soil microbial diversity manipulation experiment to explore how removing fungi from inocula impacts estimates of carbon use efficiency under a range of methodological errors. Briefly, microbial communities were extracted from temperate deciduous forest soil and either the complete (“fungi + bacteria”) or less than 0.8uM fraction (“filtered”; “bacteria only”) was used to inoculate an artificial soil matrix. This matrix consisted of 70% acid-washed sand, 20% muffled and acid-washed silt, and 10% calcium chloride-treated bentonite clay, initially amended with mixed deciduous leaf litter DOC, 2X roller media [18], VL55 minerals and yeast extract. The communities were grown for four months, with weekly additions of 0.5mg g soil^−1^ cellobiose and 0.05mg g soil^−1^ ammonium nitrate solutions as sources of C and N, respectively, for the first three months.

CUE was then measured by adding ^18^O-H_2_O to 20% of the final water present to subsamples of the soil. Samples were prepared identically, only using ^16^O-H_2_O, as controls for background heavy oxygen incorporation. The samples were then placed in sealed tubes for 24 hours and the CO_2_ produced during this time measured using an IRGA. The soil samples were stored at −80C until DNA extraction using the Qiiagen Powersoil HTP kit. The resultant DNA was quantified using PicoGreen (Invitrogen), and its ^18^O enrichment was measured using IRMS at the UC Davis Stable Isotope Facility. CUE was calculated as per [12]. The abundance of total bacteria and total fungi was assessed by real-time quantitative PCR (qPCR) using 16S rRNA primers [19] and ITS primers [20], respectively. The abundance in each soil sample was based on increasing fluorescence intensity of the SYBR Green dye during amplification. Preceding qPCR assay an inhibition test was performed by running serial dilutions of DNA extractions and no amplification inhibition was detected. The qPCR assay was carried out in a 15 ul reaction volume containing 2 ng of DNA, 7.5 ul of SYBR green (QuantiFast SYBR Green PCR Master Mix) and 1 uM of each primer. Two independent qPCR assay were performed for each gene. The qPCR effciencies for both genes ranged between 85 and 105%. 16S qPCR conditions were: 15 minutes at 95C; 40× 15s @ 94°C, 30s @ 55°C, 30s @72°C; and a melting curve. ITS qPCR conditions were: 15 minutes at 95°C; 40× 15s @ 94°C, 30s @ 46°C, 30s @ 72°C; and a melting curve. These values were corrected to a genome counts basis using median values from [21] for ITS copies and from [22] for bacterial 16S ribosomal RNA operon copy number.

### 3.3 Shiny app and theoretical sensitivity simulations

Each simulation was set up under a series of biologically-plausible scenarios. Fungal:bacterial ratio and total community size are presented in terms of DNA, as this is the unit of growth measurement for the ^18^O method. Definitions for parameters are found in Table 1, and for variables defined in the equations below in Table 2. Values without subscripts denote true MBC, DNA, and MBC:DNA ratios, while values with subscripts denote observed values were DNA (d), MBC (c), or both (dc) to be extracted inefficiently:

**Table 2:** Variables defined in the microbial biomass carbon calculations

| Parameter | Description | units |
| --- | --- | --- |
| totalDNA | The true total mass of the DNA pool in the soil | ug g<sup>-1</sup> soil |
| FBratio | Fungal fraction of total DNA pool | dimensionless |
| MBC | The true total mass of the microbial biomass carbon pool in the soil | ug g<sup>-1</sup> soil |
| MBC<sub>c</sub> | The apparent total mass of the microbial biomass carbon pool in the soil, given inefficient microbial biomass carbon extraction | ug g<sup>-1</sup> soil |
| totalDNA<sub>d</sub> | The apparent total size of the true DNA pool in the soil, given a DNA extraction inefficiency | ug g<sup>-1</sup> soil |
| MBCDNA | True microbial biomass carbon to DNA mass ratio of starting community; Used to convert the DNA growth increment into microbial biomass carbon growth. | dimensionless |
| MBCDNA<sub>c</sub> | Apparent microbial biomass carbon to DNA mass ratio of starting community, given that microbial biomass carbon is not completely extracted; Used to convert the DNA growth increment into microbial biomass carbon growth. | dimensionless |
| MBCDNA<sub>d</sub> | Apparent microbial biomass carbon to DNA mass ratio of starting community, given that DNA is not completely extracted; Used to convert the DNA growth increment into microbial biomass carbon growth. | dimensionless |
| MBCDNA<sub>dc</sub> | Apparent microbial biomass carbon to DNA mass ratio of starting community, given that both DNA and microbial biomass carbon are not completely extracted; Used to convert the DNA growth increment into microbial biomass carbon growth. | dimensionless |
| GR<sub>mean</sub> | Growth rate for simulations when bacteria and fungi are assumed to grow at the same community-level mean | day<sup>-1</sup> |
| TrueMBC<sub>same</sub> | The true new microbial biomass carbon produced during the CUE incubation, assuming bacteria and fungi grow at the same rate (GR<sub>mean</sub>) | ug C day<sup>-1</sup> |
| MBC<sub>samed</sub> | The apparent new microbial biomass carbon produced during the CUE incubation, given that not all the DNA is extracted from the growing community and assuming that bacteria and fungi grow at the same rate (GR<sub>mean</sub>) | ug C day<sup>-1</sup> |
| MBC<sub>samec</sub> | The apparent new microbial biomass carbon produced during the CUE incubation, given that not all the microbial biomass carbon is extracted from the growing community and assuming that bacteria and fungi grow at the same rate (GR<sub>mean</sub>) | ug C day<sup>-1</sup> |
| MBC<sub>samedc</sub> | The apparent new microbial biomass carbon produced during the CUE incubation, given that not all the DNA and microbial biomass carbon are extracted from the growing community and assuming that bacteria and fungi grow at the same rate (GR<sub>mean</sub>) | ug C day<sup>-1</sup> |
| DNA<sub>bias</sub> | The fraction of new oxygen in DNA which comes from extracellular water (ie the 18O-water added) rather than other sources | dimensionless |
| TrueMBC<sub>diff</sub> | The true new microbial biomass carbon produced during the CUE incubation assuming bacteria and fungi grow at different rates | ug C day<sup>-1</sup> |
| MBC<sub>diffd</sub> | The apparent new microbial biomass carbon produced during the CUE incubation assuming bacteria and fungi grow at different rates, given that not all the DNA is extracted from the growing community | ug C day<sup>-1</sup> |
| MBC<sub>diffc</sub> | The true new microbial biomass carbon produced during the CUE incubation assuming bacteria and fungi grow at different rates, given that not all the microbial biomass carbon is extracted from the growing community | ug C day<sup>-1</sup> |
| MBC<sub>diffdc</sub> | The true new microbial biomass carbon produced during the CUE incubation assuming bacteria and fungi grow at different rates, given that not all the DNA and microbial biomass carbon are extracted from the microbial community | ug C day<sup>-1</sup> |

A community of size *totalDNA* was generated as a function of the fungal fraction of the total DNA pool (*FBratio*), where 30 was used as an arbitrary multiplier to determine the amount of DNA, and 3 as an additive factor to ensure that bacteria-only (FBratio of zero) still had DNA.

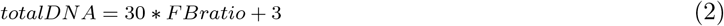

The corresponding amount of microbial biomass carbon (*MBC*) is calculated as the sum of the biomass carbon of bacteria and fungi, which are the products of their DNA and MBC:DNA ratios.

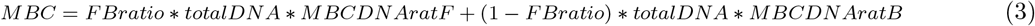

Since the MBC:DNA ratio of bacteria (*MBCDNAratB*) is generally larger in fast-growing and well-fed cells [42, 43], we also added the option to allow this ratio to vary as a function of bacterial growth rate. This was done by scaling the ratio between the minimum and maximum *MBCDNAratB* values observed in the literature over the range of bacterial growth rate (*GRbact*) values observed (Table (1)), and assuming a linear relationship:

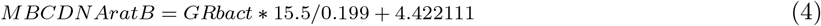

which is the solution of

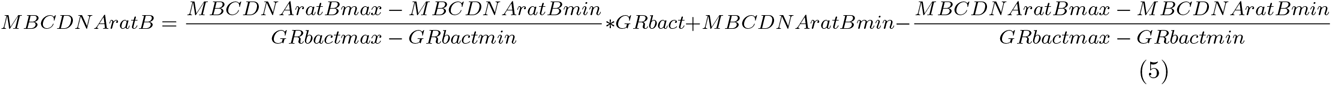

The MBC:DNA ratio of the starting community is therefore:

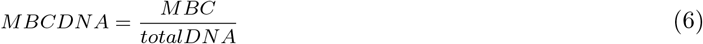

However, DNA is not completely extracted from soil microbes, with some evidence for a higher extraction efficiency for bacteria (*DNAexteffB*) than fungi (*DNAexteffF*) [31]. Spores may be extracted with even lower efficiency [54], but do not contribute to growth so do not play into our calculations. The observed total DNA observed assuming inefficient DNA extraction (*totalDNA_d_*) is then:

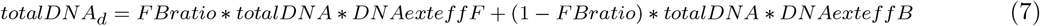

The corresponding MBC:DNA ratio assuming inefficient DNA extraction (*MBCDNA*_*d*_) is:

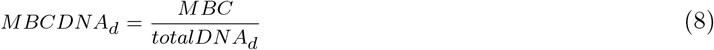

MBC is also inefficiently extracted, with chloroform fumigation extraction capturing the true fungal (*MBCexteffF*) and bacterial (*MBCexteffB*) biomass carbon present with different effciencies [48]. *MBC*_c_ represents the total amount of microbial biomass observed after accounting for this chloroform fumigation extraction inefficiency:

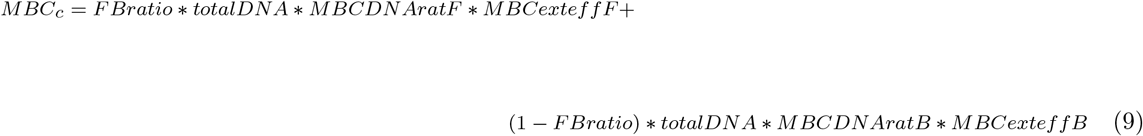

And the corresponding MBC:DNA ratio is:

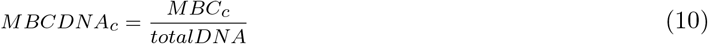

If ineffciencies in both MBC and DNA extraction must be accounted for, then the apparent MBC:DNA ratio (*MBCDNA_dc_*) is:

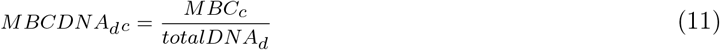

Steps 4-6 therefore show the MBC:DNA ratios a researcher converting the new DNA produced to MBC would use if they were unaware of extraction biases and did not account for differences in bacterial and fungal growth rates (below).

We assume growth during the incubation is representative of overall community growth. In other words, the community is assumed to be in a steady state and the rate of turnover of a given taxon matches its growth. In turn, the turnover of DNA in the environment is proportionate to its abundance [55]. If bacteria and fungi grow at the same rate, then the community-level growth rate (GRmean) can be set to vary as a function of the community composition:

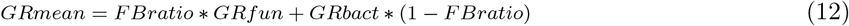

The corresponding true increase in MBC for bacteria and fungi when they are assumed to grow at the same rate (*TrueMBCsame*) is:

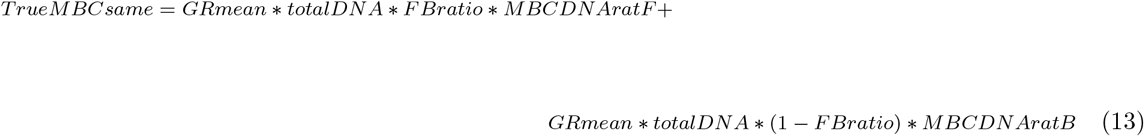

However, we may not “see” all this growth because extracellular water is not the sole source of oxygen in DNA. Rather, anywhere from 4-70% of oxygen in DNA may come from metabolic water [27, 26, 28]. We refer to this bias towards using extracellular rather than intracellular water as *DNAbias*, which is the fraction of DNA oxygen derived from extracellular water.

Subsequently, if just DNA is extracted inefficiently then the corresponding apparent new MBC produced (MBCsame_d_) is:

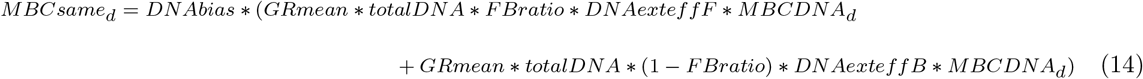

If just MBC is extracted inefficiently, then the corresponding apparent new MBC produced (MBCsame_c_) is:

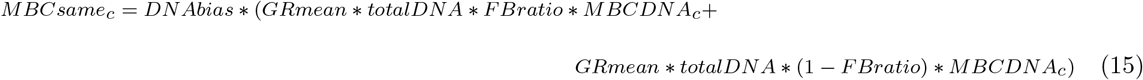

If both MBC and DNA are extracted inefficiently, then the apparent new MBC produced (MBCsame_dc_) is:

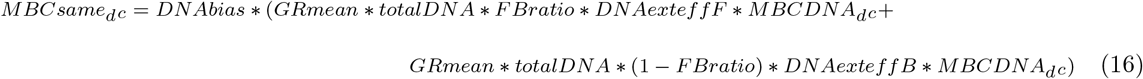

When bacterial growth rate (*GRbact*) and fungal growth rate (*GRfun*) differ, the true MBC produced (*TrueM-BCdif*) is:

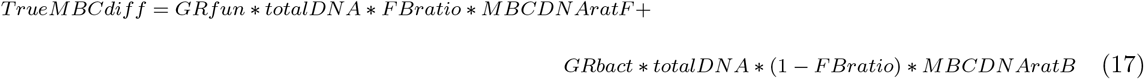

And the values for the true MBC produced under the various extraction bias scenarios are:

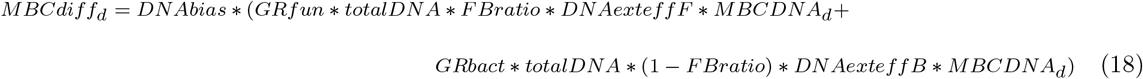

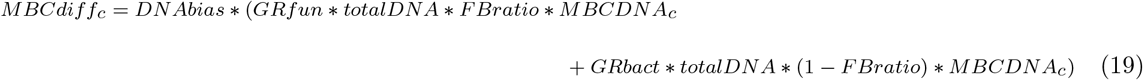

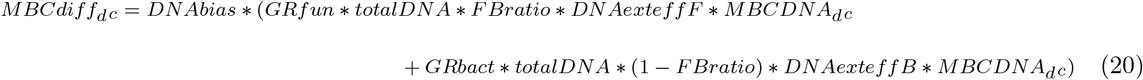

One of the assumptions of the ^18^O-CUE method is that the turnover of labeled biomass is negligible over the course of the incubation. Assuming a steady microbial community biomass, the corresponding bulk turnover rates of 0.3 to 7% per day above indicate that this expectation is reasonable. However, the true DNA growth rate is likely to be higher, and the impact on estimates of microbial carbon growth to be mixed. Dormancy estimations vary widely, from 6 to 96% of the community observed as dormant [25, 23]. Considering a scenario in which 96% of the community is dormant leads to a 24-fold underestimation of growth rate (0.96/(1-0.96)) due to dilution with the bulk pool, but only minimal underestimation if 6% are. Furthermore, factors such as predation could decrease apparent growth rate through the inefficient re-allocation of labeled nucleic acids from primary to secondary consumers, particularly if predators selectively consume community members [56] within a narrow size range [57]. Finally, as a result of the “live fast, die young” adage often attributed to copiotrophs, CUE is likely to be particularly underestimated when growth is concentrated in a small but rapidly growing fraction of the population compared to a larger but slower growing fraction. Our simulations accounted for an active community fraction varying from 1 to 99

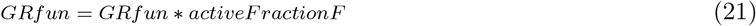

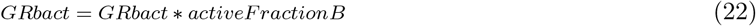

### 3.4 Sensitivity of CUE to fungal removal

We assessed the sensitivity of observed CUE to various methodological assumptions using a few modifications to account for observed fungal:bacterial ratio. Unlike the simulations above, we wished to retain the inter-sample differences in MBC:DNA ratio and growth. Therefore we applied modifying factors to the original data using expected ratios between bacterial and fungal parameters, rather than imposing fixed values for these organism classes as above.

First, we converted the observed fungal:bacterial DNA ratio based on qPCR to a F:B DNA ratio. To do this, we assumed 82 ITS copies per genome (*ITSpergenome*) (Table 1) and a median genome size of 5×10^8^ bp for fungi [21, 58], and 2.25 16S copies per genome (*16Spergenome*) and a genome size of 5×10^6^bp for bacteria [22, 58]. To get the true fungal:bacterial DNA ratio *FBratio*, we had to back-calculate from the observed ITS copies *copiesITS* and 16S ribosomal RNA copies *copies16S* from qPCR. We then accounted for ineffciencies in DNA extraction as follows:

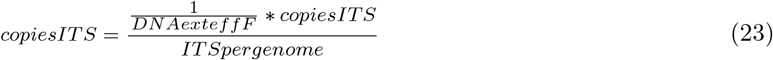

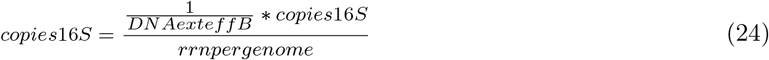

And then convert the 16S and ITS copies to fungal (*FDNA*) and bacterial (*BDNA*) DNA mass per gram of soil as follows:

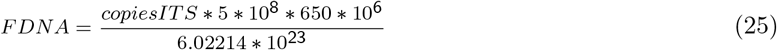

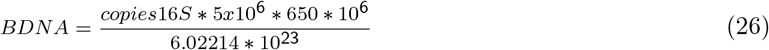

Where 650*10^6^ is the molecular weight of the average DNA basepair in *µ*g and 6.02214*10^23^ is Avogadro’s constant. So the corresponding extraction-efficiency and marker gene per genome base pair corrected fungus DNA: bacteria DNA ratio (*FBratio*) is:

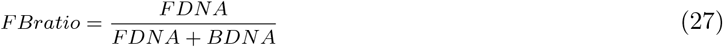

The corresponding corrected total DNA (*totalDNAActual*) in the initial pool (active and inactive) used for MBC:DNA ratio calculation is:

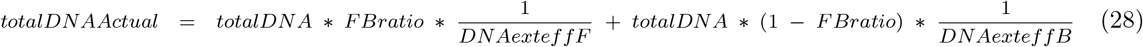

We can then calculate relative fungal and bacterial contributions to the MBC pool for fungi (*fcont*) and bacteria (*bcont*), the actual amount of MBC (*MBCactual*) and the MBC:DNA ratios for each group as follows:

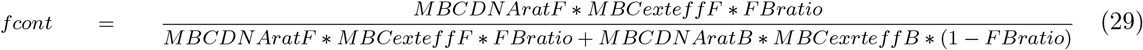

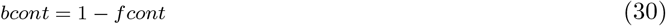

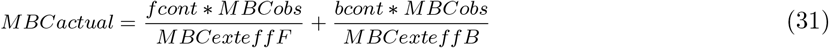

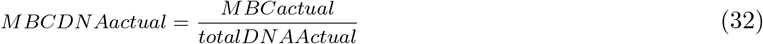

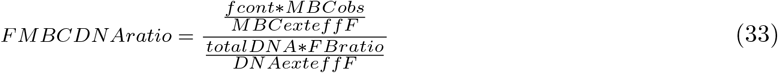

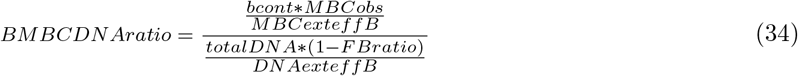

Now we calculate the *FBratio_active*, which is the fraction of new growth attributed to fungi during the incubation. It is a function of the relative growth rates of bacteria and fungi, as well as their FBratio in the starting bulk community and the fraction of the cells which are active, rather than dormant.

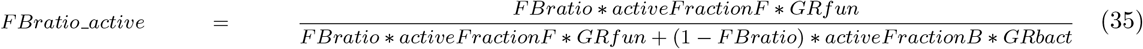

We can then account for DNA extraction (in)efficiency and the use of intracellular water/other sources of DNA oxygen for the growing community:

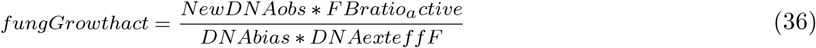

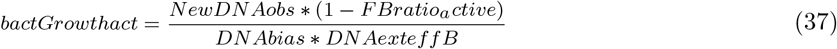

Finally, we convert these DNA growth to the MBC growth which occurred after applying our methodological bias corrections:

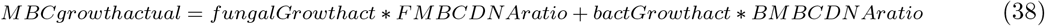

And calculate CUE using the observed respiration rate (per day):

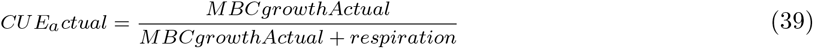

## 4 Results and Discussion

### 4.1 Growth bias depends on extraction bias and FB ratio

Our Shiny app simulations showed that observed microbial growth deviated most from true microbial growth at intermediate fungal:bacterial ratios, and when fungi and bacteria grew at different rates (Figure 2). If only bacteria or fungi are present the active community is better represented by the total community and the MBC:DNA ratio of the active community is as well represented as possible in the MBC:DNA ratio of the starting community. We also found that if groups of microbes with distinct MBC:DNA ratios grow at the same growth rate, then the ability of the ^18^O method to reliably estimate the increase in MBC is insensitive to any differences in the DNA extraction efficiency. This is demonstrated as the observed and actual MBC growth falling on the 1:1 line over all FB ratios, and can be explained by the DNA being underestimated by equivalent amounts in both the total community used for MBC:DNA conversion and the active community extracted. However, mis-estimating the MBC extraction efficiency leads to incorrect microbial growth values whether or not bacteria and fungi grow at the same rate (Figure 2, center column). While a mathematically simple scenario to explain, this is particularly alarming because CFE extraction efficiency depends on a wide range of experimentally-relevant features. This includes the ratio of intracellular (cytoplasm) to extracellular (membrane, extracellular polysaccharides, proteins) carbon, which is known to differ with community structure and growth rate [48, 50] and edaphic parameters such as soil pH, water and clay contents [59, 60].

**Figure 2:**
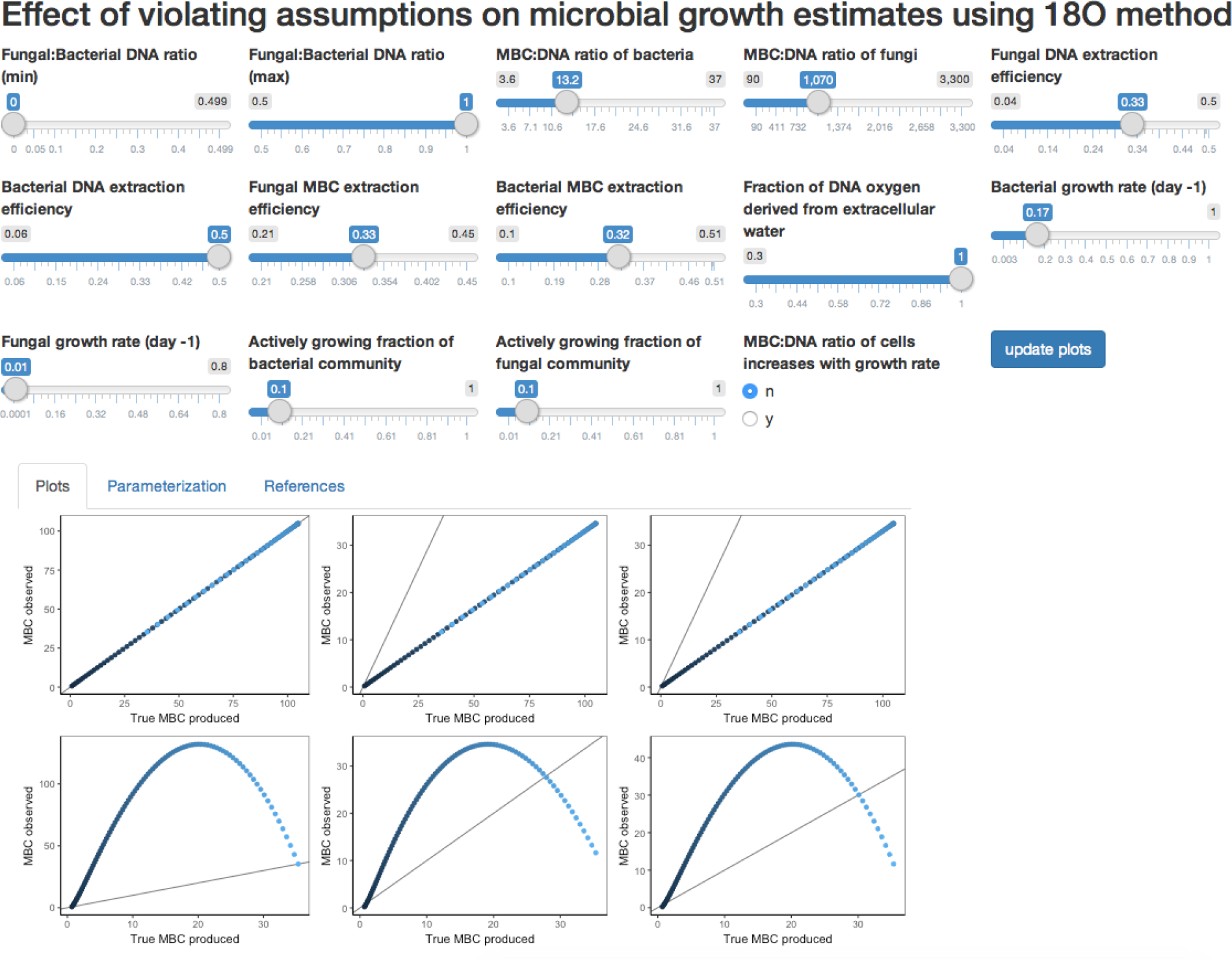
Screenshot of Shiny app used to visualize the effect of methodological error on microbial biomass carbon estimates. Each point denotes a community simulated for a different fungal:bacterial DNA ratio, with darker points representing a more bacterial community (in this instance, FB = 0) and lighter blue a more fungally-dominated community (here, FB ratio of 1). The black diagonal denotes the 1:1 line, such that values above the line indicate overstimation of biomass, and those below indicate underestimation. Top row: bacteria and fungi grow at the same rate. Bottom row: bacteria and fungi grow at distinct rates. Left column: DNA extracted inefficiently. Center column: MBC extracted inefficiently. Right column: MBC and DNA both extracted inefficiently.

### 4.2 CUE estimates are sensitive in the presence of metabolic water

Using a sensitivity analysis and varying one factor at a time, we found that the deviation of observed microbial growth from true growth was most sensitive to metabolic water content across all extraction scenarios (Figure 3); the sensitivity value was 1 throughout. The first uses of the ^18^O method for CUE assumed that all DNA oxygen came from extracellular water [12], but it is known that *E. coli* only derives around 30% of its DNA oxygen from intracellular water when grown on rich media in the lab [26, 29]. Under the less-than ideal conditions in the soil, the value is likely to be higher - from 70-98% of oxygen from extracellular water - as the contribution of intracellular water to DNA oxygen is lower in slower growing *E. coli* [26] and *B. subtilis* [28]. The degree of ^18^O enrichment in the phosphate backbone also decreases with temperature [61]; since growth rate often increases with temperature, this is another mechanism by which growth may be underestimated in the fastest growing communities. On the other hand, the recycling of nucleotides and “cryptic growth” may be more important in slow-growing and nutrient-starved organisms, preferentially hiding growth in these communities. Moreover, we observed the suppression of respiration after addition of ^18^O H_2_O compared to ^16^O water in three temperate deciduous forest soils (Figure S6), which could indicate that ^18^O-H_2_O undersestimates growth by suppressing metabolism. We still lack precise estimates of how important non-extracellular water sources are for DNA oxygen under *in-situ* conditions for bacteria or any conditions for fungi, making them important areas for future research.

**Figure 3:**
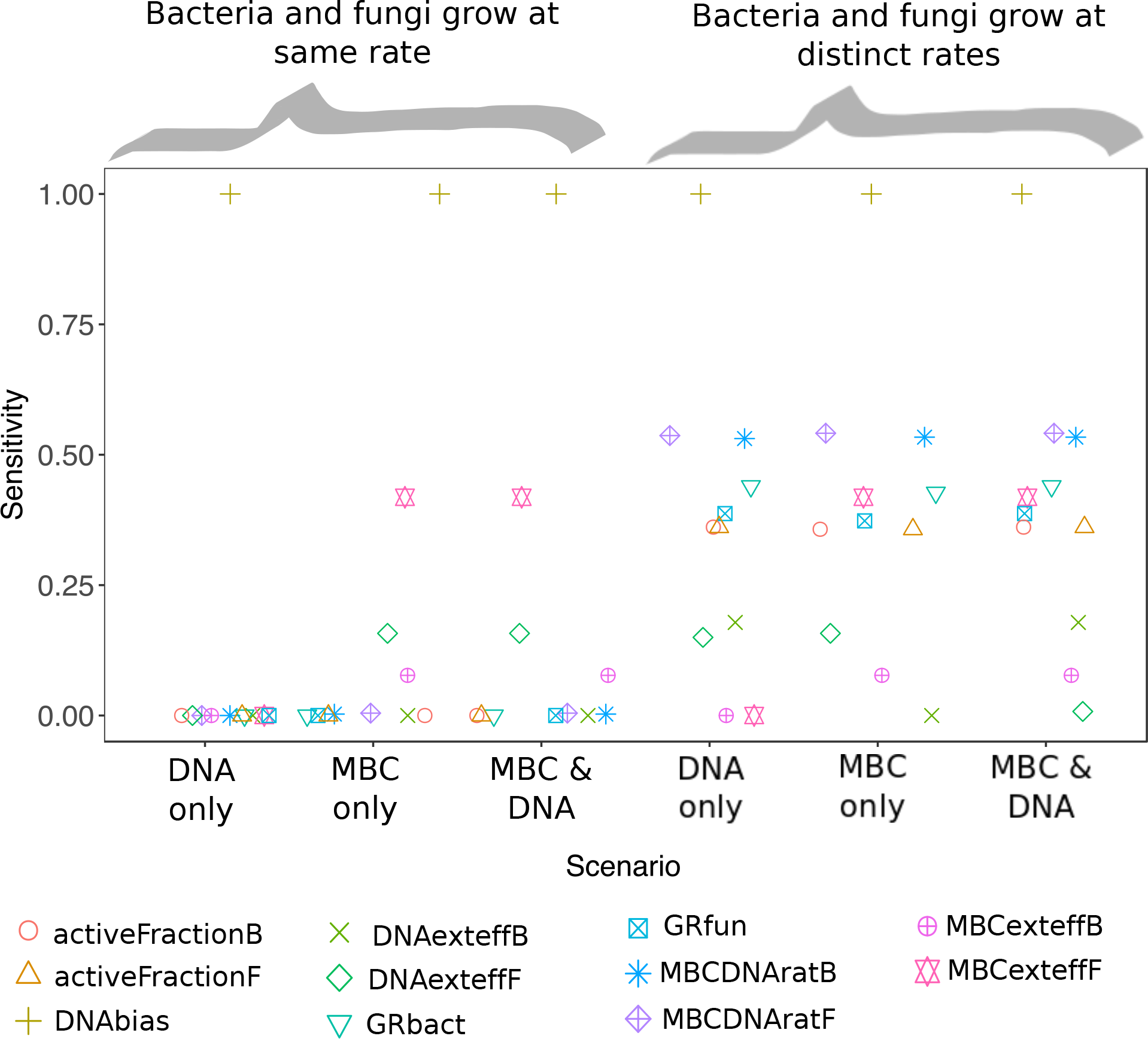
sensitivity of difference between true and observed microbial growth values to violating various assumptions. Left: bacteria and fungi grow at different rates; right both grow at the population level mean.

**Figure 4:**
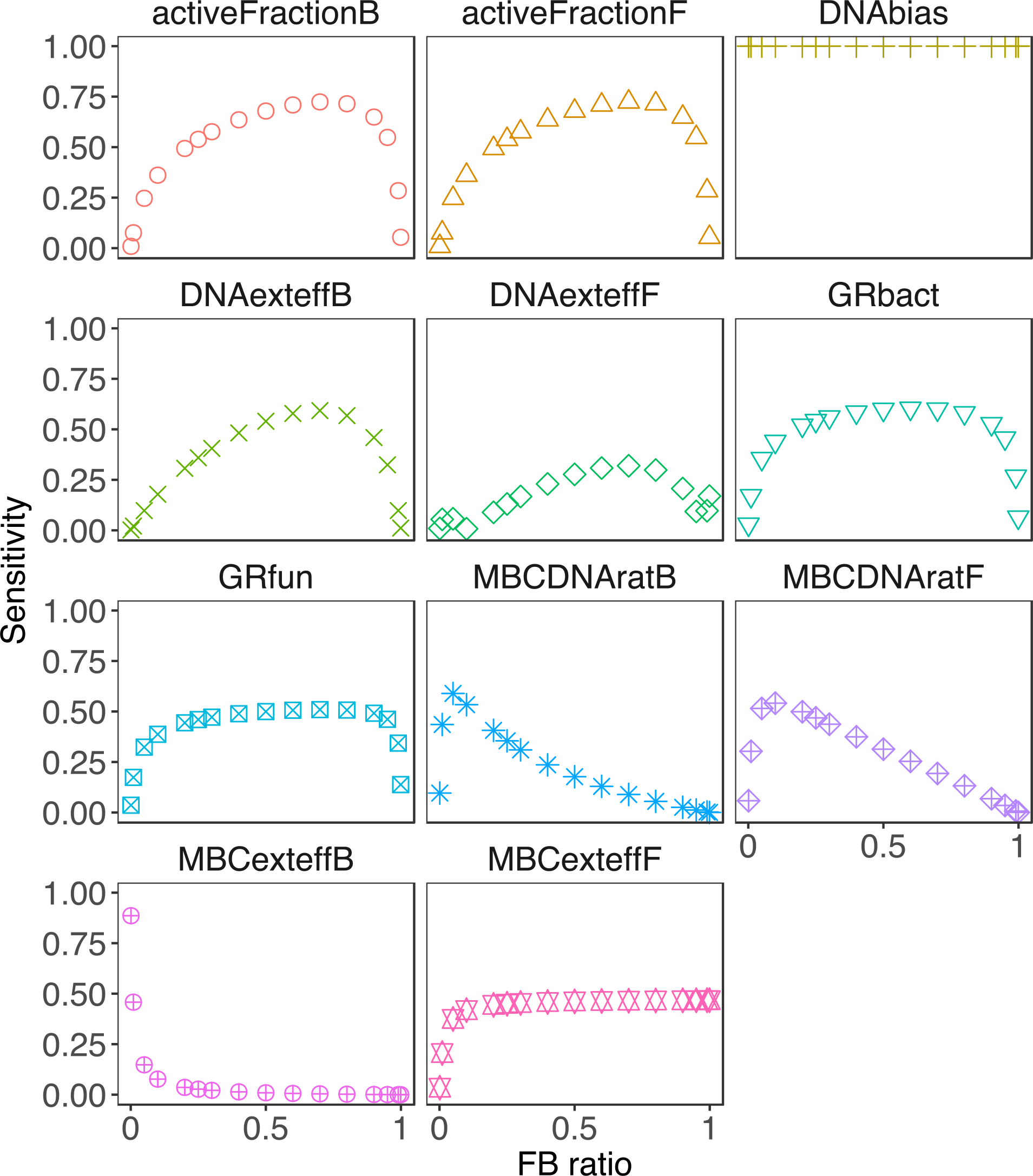
sensitivity of MBC growth estimate error to variation in biological parameters and methodological errors. The plotted scenario assumes that DNA and MBC are both under-extracted, and that F and B grow at different rates. Results for the remaining scenarios in (Figure 3) can be found in figures S1-S5. Parameters are defined in Table 1

### 4.3 Sensitivity of growth to methodological bias depends on heterogeneity in growth rates

With the exception of intracellular water contribution, MBC sensitivity to changes in the other parameters depended on both assumed extraction biases and whether bacteria and fungi grew at same or different rates (Figure 3). In general, estimates were more sensitive to changes in the parameters when bacteria and fungi grew at different rates rather than some community-level mean, in large part because the MBC:DNA ratio observed for the whole community was no longer representative of the growing population. Fungal MBC:DNA extraction efficiency had a similar effect on how far expected growth deviated from observed growth independent of whether bacteria and fungi grew at the same rate. Errors were also sensitive to bacterial MBC extraction efficiency, but less so. This is because despite slow DNA-based growth in the baseline condition, fungi have a very large MBC:DNA ratio and so contribute disproportionately to the MBC estimate. In a similar thread, errors were less sensitive to fungal DNA extraction efficiency because their growth rate is minimal under baseline conditions. To address this slow growth, researchers sometimes add different amounts of 18O-water to soils or incubate for different periods based on the growth rates of soils [5].

### 4.4 Sensitivity of errors in MBC estimations depend on FBratio

Fungal to bacterial DNA ratio affected which parameters MBC estimates were most sensitive to, with these sensitivities also differing in their sensitivity to F:B ratio. For instance, MBC error had a sensitivity of approximately 0.5 over all intermediate values of fungal and bacterial growth rate, but decreased precipitously towards zero at FB ratios approaching 0 or 1. By contrast, sensitivity to bacterial DNA extraction efficiency was greatest around a FB ratio of 0.7, and to fungal DNA extraction efficiency at an FB ratio of 0.1-0.2. Sensitivity of MBC extraction efficiency for bacteria was almost zero, except at F:B ratios below 0.1, but almost 1 for fungal MBC extraction efficiency under these same scenarios. Assuming the DNA content and rrN/ITS copy numbers in table 2, the F:B DNA ratio in both soil metagenomic sequences [62] and qpcr [63, 22] datasets are often less than 10%. As such, many soil samples are within the range of F:B DNA ratios where deviations in CUE estimates are highly sensitive to even small changes in fungal dominance, such that related samples within a study may differ in the kinds of methodological assumptions they are most sensitive to.

### 4.5 Effect of fungal removal on CUE

Fungal:bacterial ratio is one of the oldest and coarsest ways of differentiating microbial communities, with the ratio typically decreasing with depth and increasing with carbon content [64, 63]. We found that our conclusions achieved with our simulations regarding the effects of fungal removal on CUE were confirmed by empirically excluding fungi from an artificial soil inocula. This is despite the observation that microcosms with bacteria only or both bacteria and fungi differed in the parameters they were most sensitive to (Figure 6). For instance, bacteria only microcosms were 2.9× more sensitive to bacterial biomass carbon extraction efficiency, but less than 1% as sensitive to dormancy than were the microcosms with both bacteria and fungi. This led to CUE estimates responding differently to the same assumption in the two community types (Figure 5). The observed, “uncorrected” CUE was on average only 25% as high in communities with fungi excluded than those with fungi for the raw data (Figure 5, solid grey line). No scenarios led CUE in bacteria-only microcosms to approach that of the mixed fungal and bacterial microcosms (ratio = 1; dashed line). This is likely because bacteria dominated in both bacteria-only and mixed microcosms, and was estimated to account for greater than 99% of the DNA. This imbalance in fungal abundance is much greater than the MBC:DNA ratio for fungi would need to be in order to overcome their much slower growth rates compared to bacteria in our simulations.

**Figure 5:**
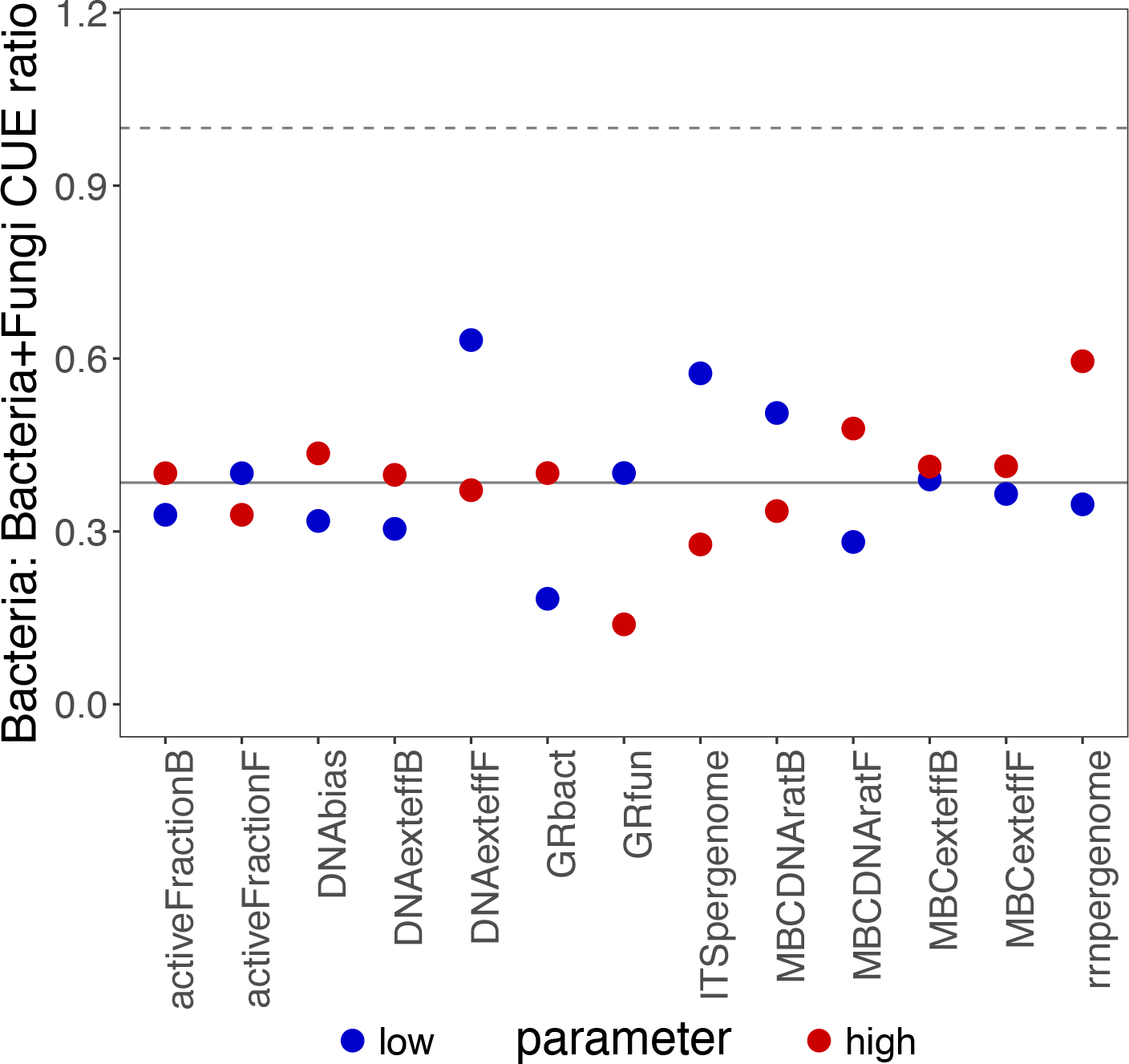
Ratio of carbon use efficiency in artificial soil microcosms inoculated with the “bacterial” (≤0.8um) fraction of soil microbial communities to the value in “complete” soil communities (“bacteria and fungi”). The x-axis denotes which one of the parameters was tested, and dot colour denotes whether the simulated CUE correction was applied at the highest value observed in a literature search, or the lowest. Each value is the median ratio for 6-8 raw replicates. The grey line denotes the median uncorrected CUE.

**Figure 6:**
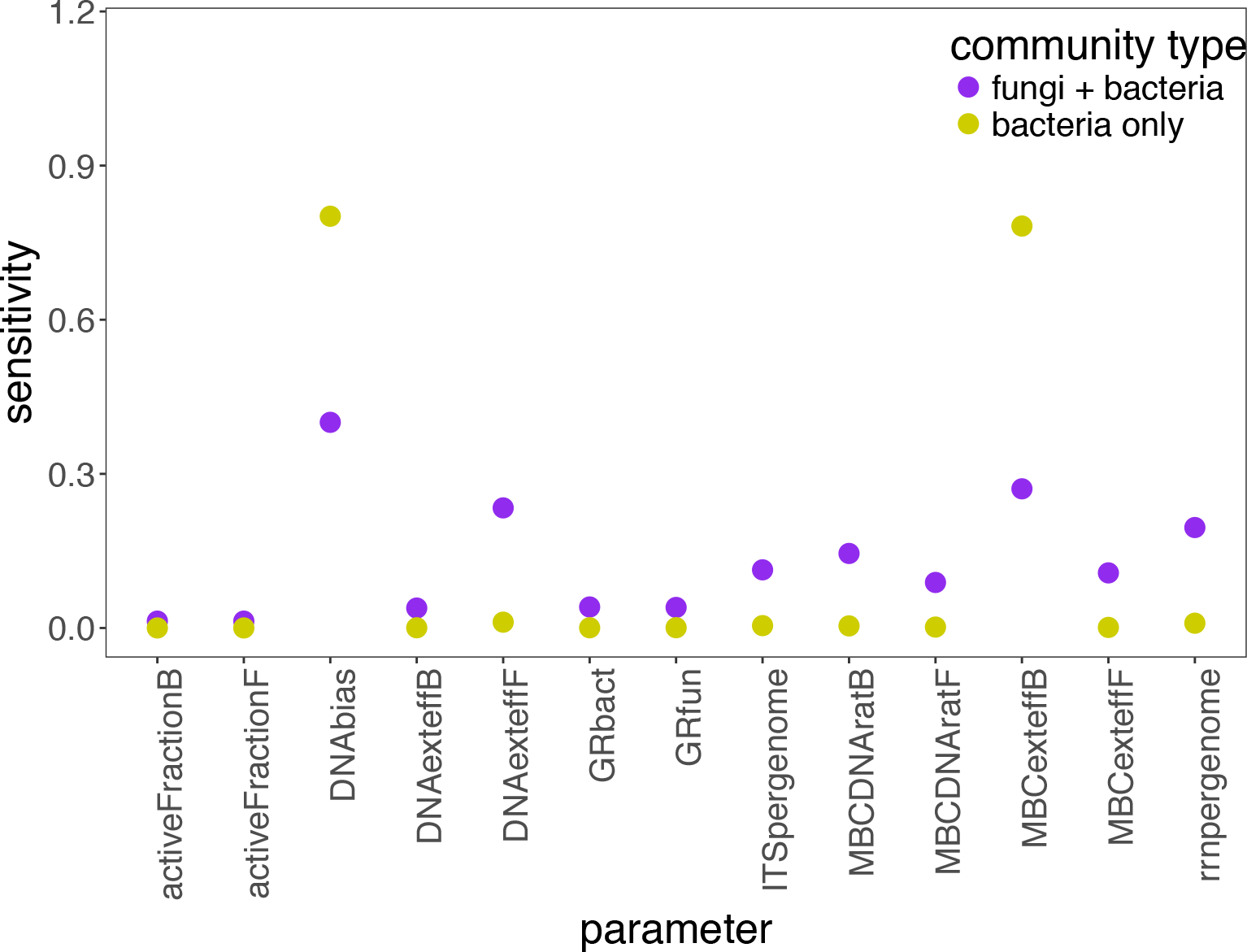
Sensitivity analysis of CUE under various methodological biases for microcosms inoculated with a filtered (“bacteria only”) or unfiltered (“bacteria and fungi”) soil inoculum. Here, sensitivity represents the deviation between the observed and simulated CUE values under the high and low parameter values presented in Table 1

Increasing the mean bacterial rrN per genome or decreasing the mean fungal ITS copies per genome increased the ratio of bacteria to bacteria + fungi CUE by decreasing the bacterial contribution to the total DNA pool. However, because fungal DNA was either absent or nearly absent from bacteria-only microcosms, this impacted the mixed community microcosms much more strongly. Since bacteria grow much faster than fungi by default in the model, reducing bacterial contribution to growth in the mixed microcosms enabled the high MBC:DNA ratio fungi to contribute more, and, in turn, increasing estimated MBC increment and CUE. All together, these results indicate that the observation of reduced CUE in communities where fungi were filtered out is not due to a single methodological bias.

Early studies of bacterial vs. fungal CUE proposed that bacteria should be less effcient than fungi because of their lower biomass CN ratio. Our results do not dispute that bacteria are less effcient than fungi, as the pattern held even with extreme corrections to CUE. Furthermore, recent work explicitly accounting for differences in bacterial and fungal growth rates and biomasses have found lower CUE in fungal-dominated communities [65]. However, our results do illustrate the benefits of sample-specific conversion factors. Microcosms differed in the assumptions their CUE estimates were most sensitive to (Figure 6), and biological differences between samples can alter the degree of methodological correction required. In other words, there are a number of possible methodological biases introduced by the act of using a single set of conversion factors for communities with and without fungi. First is that the bacterial communities are dissimilar in composition between the two treatments, with more Gram-positive Actinobacteria in bacteria-only microcosms than bacteria+fungi microcosms. The MBC:DNA ratio of the bacteria-only microcosms was very low, sometimes below 1, indicating that the bacterial biomass carbon was not efficiently extracted. By contrast, observed MBC:DNA ratios of natural soil communities generally fall between 3 and 60 [9, 45], with values as low as 3.6 for bacteria and as high as 3300 for filamentous fungi in the lab (File S1). In addition, the true MBC:DNA ratio of bacteria is lower for small, slow-growing and starving or oligotrophic cells [42, 43, 66]. Small cells have a large amount of membrane (which CFE does not effectively capture [48, 50]) relative to cytoplasm (which it does), therefore exacerbating the genuinely lower MBC:DNA ratio. Although we lack empirical evidence for smaller bacterial cells in the absence of fungi, this could explain the apparently low MBC:DNA ratio and necessitate using different extraction effciencies and MBC:DNA conversions in the two communities. However, we also note that the ^18^O method of microbial growth determination already requires a number of assumptions to be made, so making additional assumptions should be done with care.

### 4.6 Shortcomings

Many of the values used to parameterize these simulations are based on isolates grown in the lab under ideal conditions. However, microbes are known to grow very differently in the lab compared to in soil. For instance, well-fed bacterial cultures will have lower dormancy and less starvation-induced reductive cell division than those found in soil [66]. Cultivation bias towards fast-growing organisms only exacerbates this, as the (CFE-measureable) cytoplasm:(CFE-ignored) cell membrane ratio will be greater in the copiotrophic organisms we tend to study in the lab [40]. The DNA:MBC ratio has been observed to be higher in small, slow-growing cells in communities extracted from soil [42], but to remain constant over a wide range of growth rates in *E. coli* [67, 68]. Given how poorly-defined this relationship is, we did not include it as a component in our simulations, although the sensitivity of biomass increment estimates to this parameter indicate that - should such a pattern exist - it should have been accounted for. Furthermore, we note that the contribution of intracellular water to DNA oxygen was 70% for fast compared to 4% in slow-growing bacterial culture on rich media. Therefore, it is likely important to account for intersample differences in the contribution of ^18^O-H_2_O to DNA water as a function of growth rate. However, in the absence of knowledge about where bacterial and fungal growth in soil fit on this intracellular water spectrum, we did not include this parameter in our simulations. Finally, determining the true contribution of different groups of microbes to the soil DNA pool remains challenging; accurate predictions based on metagenomes are limited by both database biases and the abundance of non-coding DNA in eukaryote genomes, while imperfect primers and differences in ribosomal RNA operon copy number limit the utility of QPCR. As such, correction factors for microbial biomass carbon estimates will always be limited by the accuracy of fungal to bacterial ratios in the present simulation framework.

## 5 Conclusion

CUE is an essential descriptor of soil carbon cycling, with interesting ramifications for both the ecology and biogeochemistry of soil. There is great interest in measuring this parameter, but also a growing awareness of the various shortcomings in its quantification. Here we focused on one method - ^18^O water incorporation into DNA, arguably the most reproducible [5] - to examine how assumptions about what it actually measures affects the conclusions drawn from its estimation. We evaluated how inefficient biomolecule extraction, deviations in microbial growth rate from the population mean, and heterogeneity in microbial community composition affected how far off observed microbial growth values are from their true values. We found that measurements are particularly sensitive to the use of oxygen sources other than extracellular water, a value which has been shown to change with experimental variables such as temperature and growth rate under controlled conditions in the lab. Despite this and other possible biases affecting the CUE we observed in our lab study, our conclusions regarding reduced CUE following fungal removal held. Nonetheless, our results do not account for the possibility that the biology underlying the observed differences in CUE may necessitate sample-specific correction factors, for instance assuming lower MBC extraction effciencies in clay-rich or nutrient-poor soils compared to more organic soils. However, a more complete understanding of the constraints on and biological factors driving the importance of the biases proposed here is needed if these sample-specific correction factors are to improve - rather than worsen - the degree of measurement bias in CUE. For the time being, we therefore strongly encourage other studies to use the model script that we developed here as a springboard for evaluating how robust their own conclusions are to the various ^18^O-H_2_O CUE method assumptions.

## Supporting information

supplementary file S3 - R script

supplementary file S4 - R script

## 6 Data availability

All data and scripts used to generate the figures in this paper are provided in the supplement.

## 7 Acknowledgements

Funding for this project was provided by the Department of Energy grant DE-SC0016590 to KMD, and an American Association of University Women American Dissertation fellowship to GP. We would also like to thank Eric Morrison for fungal MBC:DNA ratio data, and to Kevin Geyer and Xiaojun Liu for conversations about the ^18^O-water method.

## 9 Supplementary files and figures

**Supplementary file S1** - review of current knowledge on pathways for water oxygen incorporation into DNA and isotopic fractionation.

**Supplementary file S2** - (app.R) Shiny app designed to visualize deviations in apparent microbial growth from the true underlying growth used to generate the data under six different scenarios. The scenarios are where fungi and bacteria show the same community level mean growth rate (top row), or distinct rates (bottom row), and, from left to right, with only DNA extracted inefficiently, just MBC extracted inefficiently, and DNA and microbial biomass carbon both extracted inefficiently. Points are coloured by fungal:bacterial ratio, with light blue at the highest FB ratio simulated, and black for the lowest (most bacterially-dominated) community. The results from the lit search used to parameterize the simulations can be found in the “Parameterization” tab, and corresponding references in the “References” tab.

**Supplementary file S3** (FileS3 190207 sensitivityCUE meanRange.R) R script used to generate data for Figure 3 and figure 4.

**Supplementary file S4** (FileS4 MicrocosmSensitivitySimulations.R) R script used to generate CUE corrections for Figure 5 and accompanying discussion.

**Figure S1.**
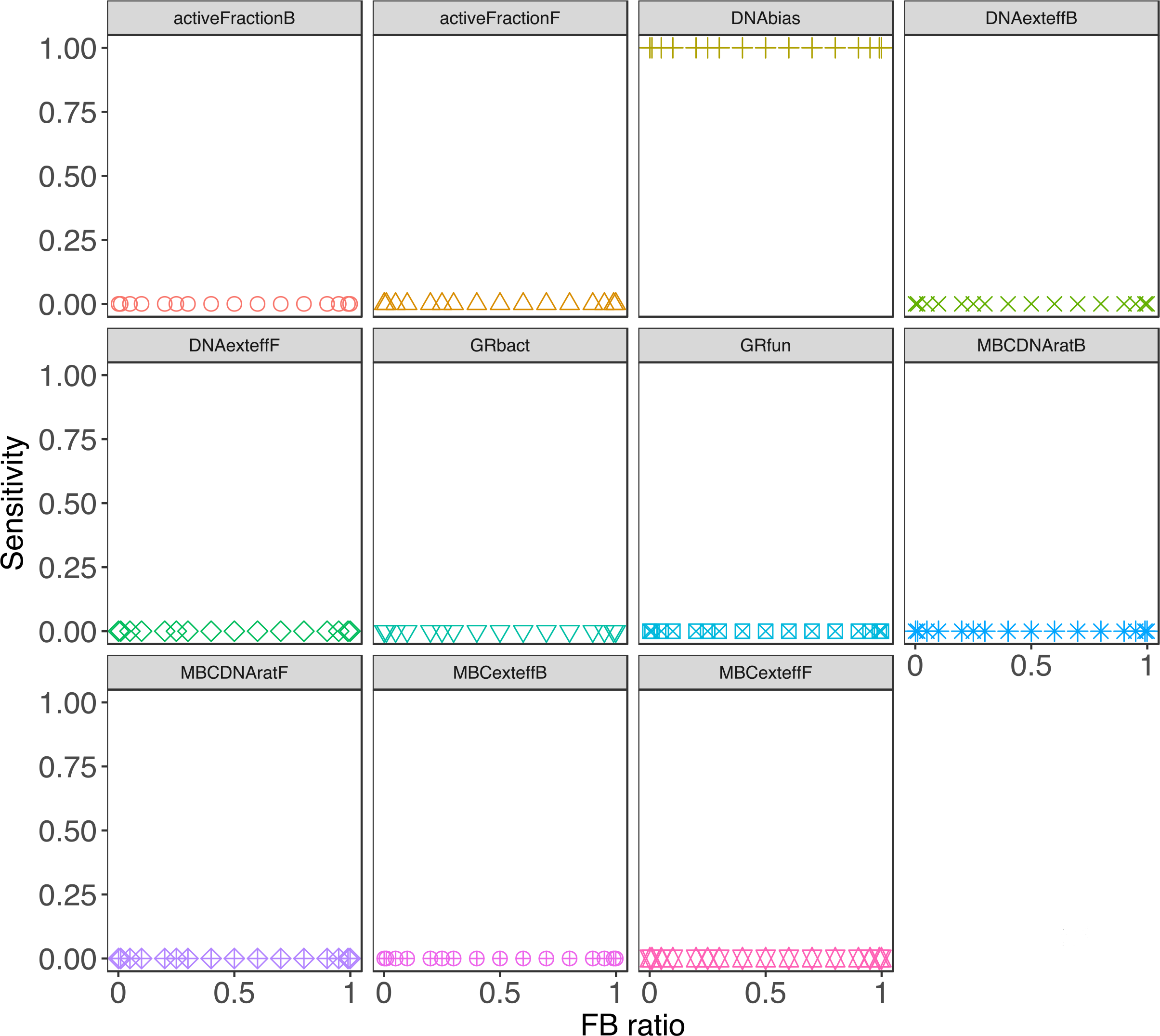
Effect of changing FB DNA ratio on sensitivity of errors in microbial growth estimates when fungi and bacteria grow at the same community level mean and only DNA extraction is incomplete.

**Figure S2.**
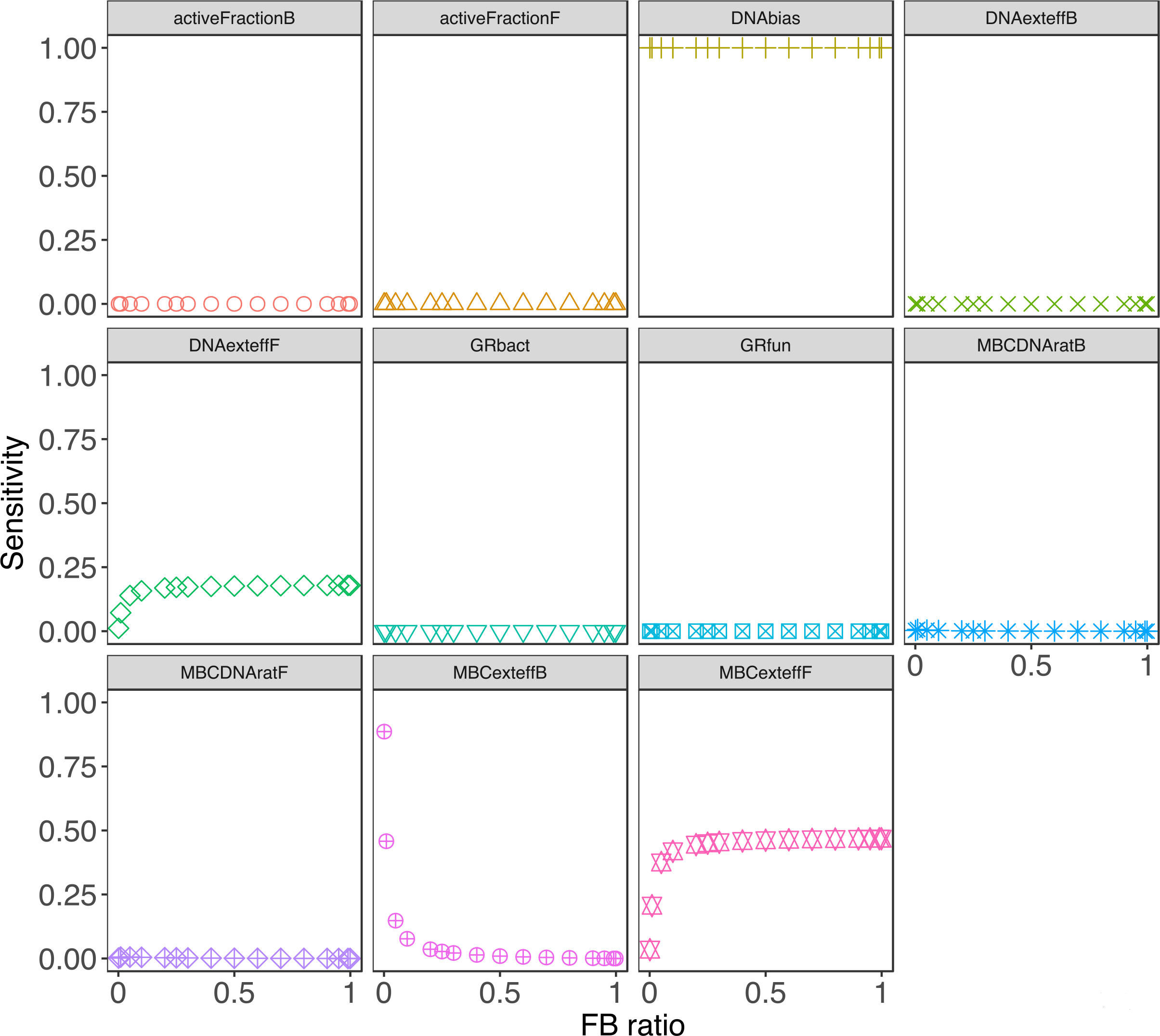
Effect of changing FB DNA ratio on sensitivity of errors in microbial growth estimates when fungi and bacteria grow at the same community level mean and only MBC extraction is incomplete.

**Figure S3.**
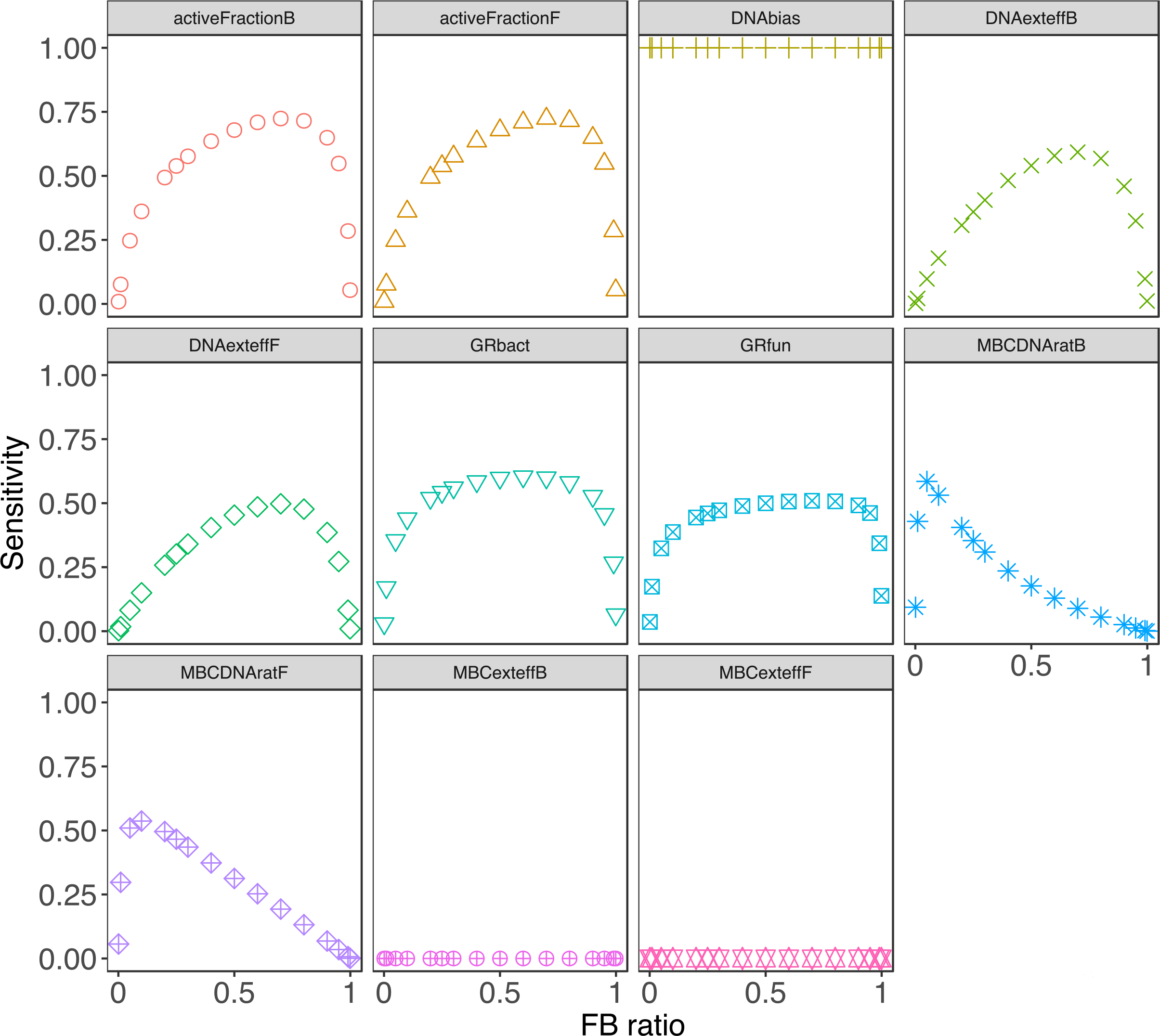
Effect of changing FB DNA ratio on sensitivity of errors in microbial growth estimates when fungi and bacteria grow at the same community level mean and both DNA and MBC extraction are incomplete.

**Figure S4.**
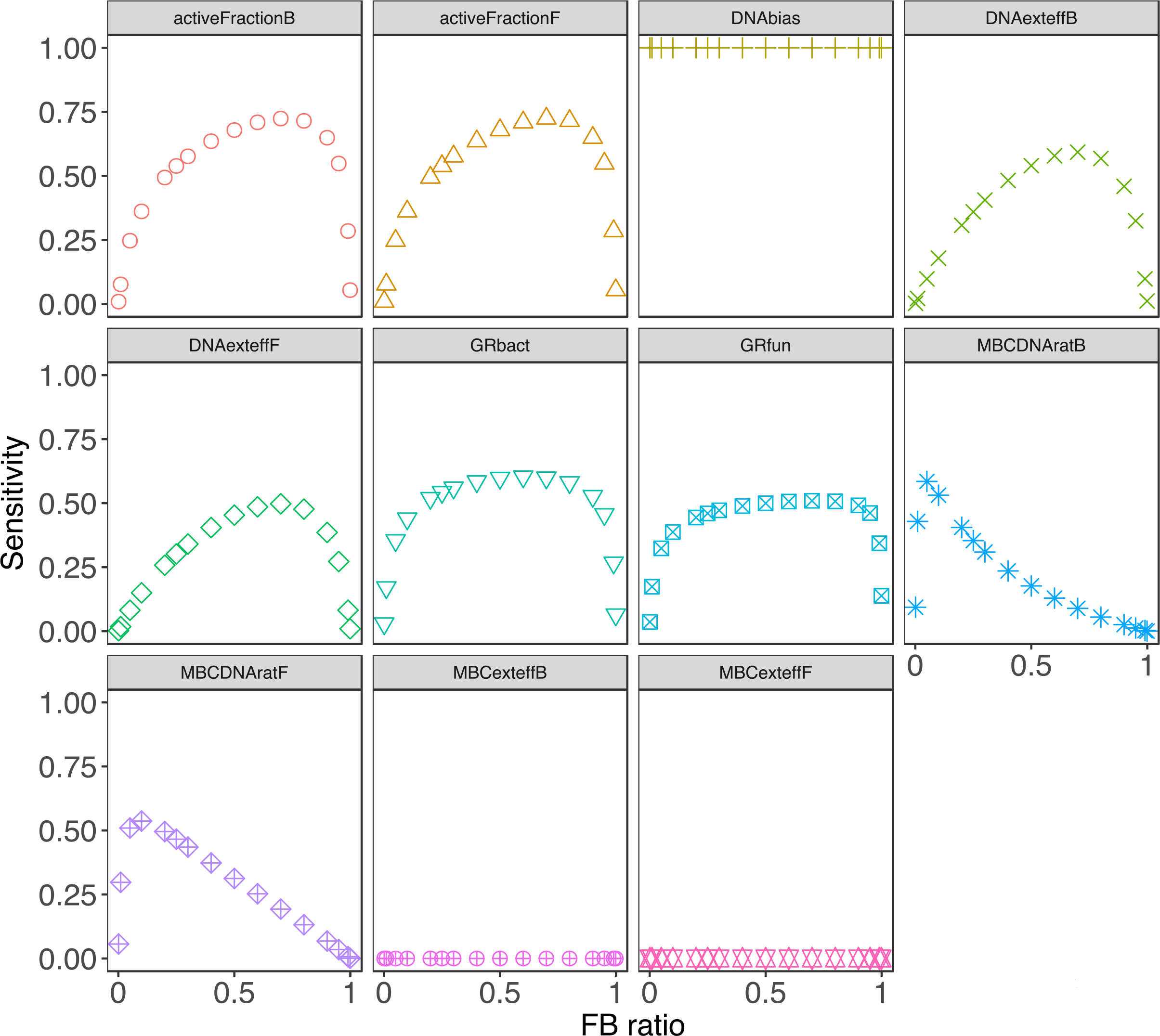
Effect of changing FB DNA ratio on sensitivity of errors in microbial growth estimates when fungi and bacteria grow at dissimilar rates and only DNA extraction is incomplete.

**Figure S5.**
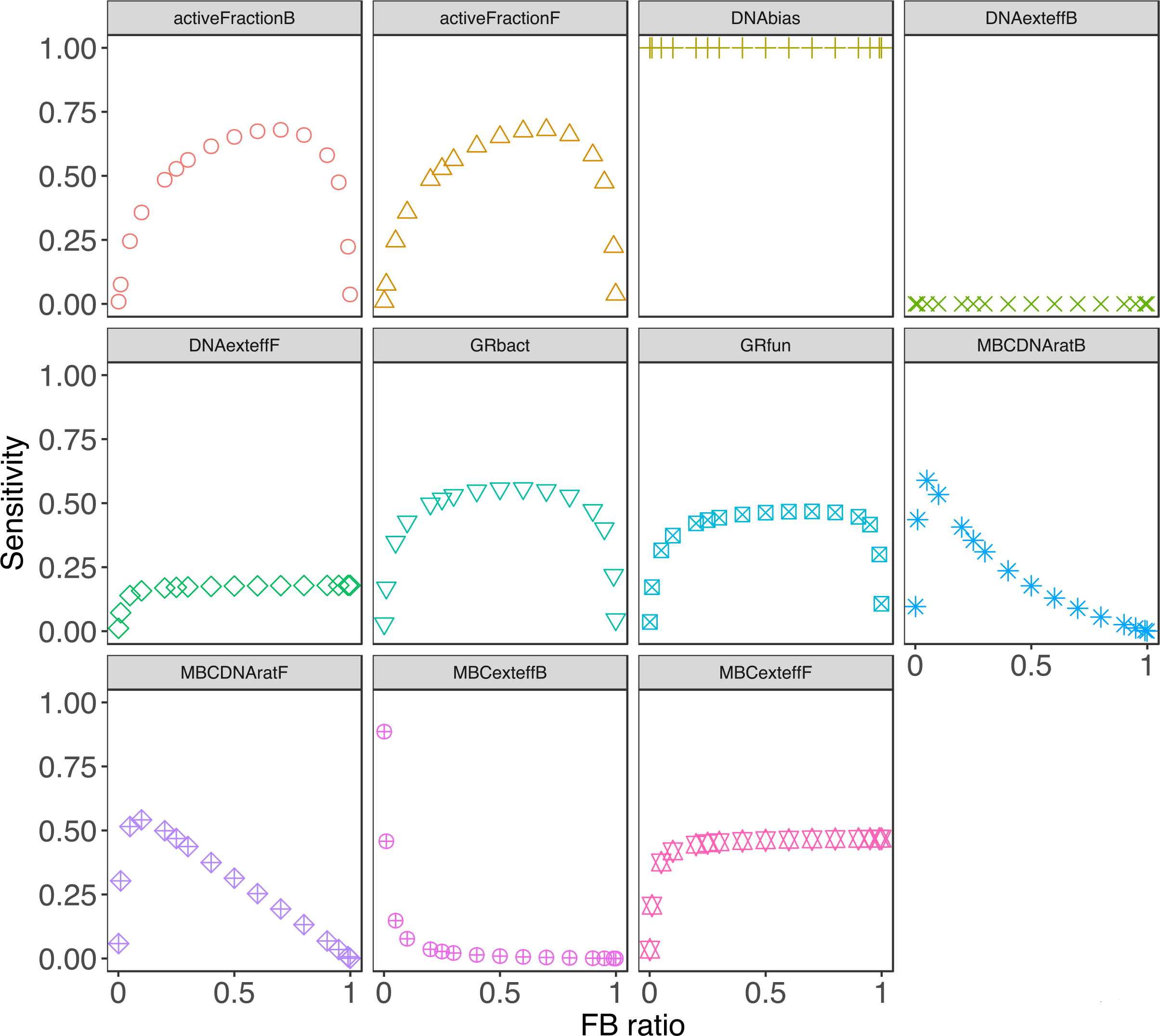
Effect of changing FB DNA ratio on sensitivity of errors in microbial growth estimates when fungi and bacteria grow at dissimilar rates and only DNA extraction is incomplete

**Figure S6.**
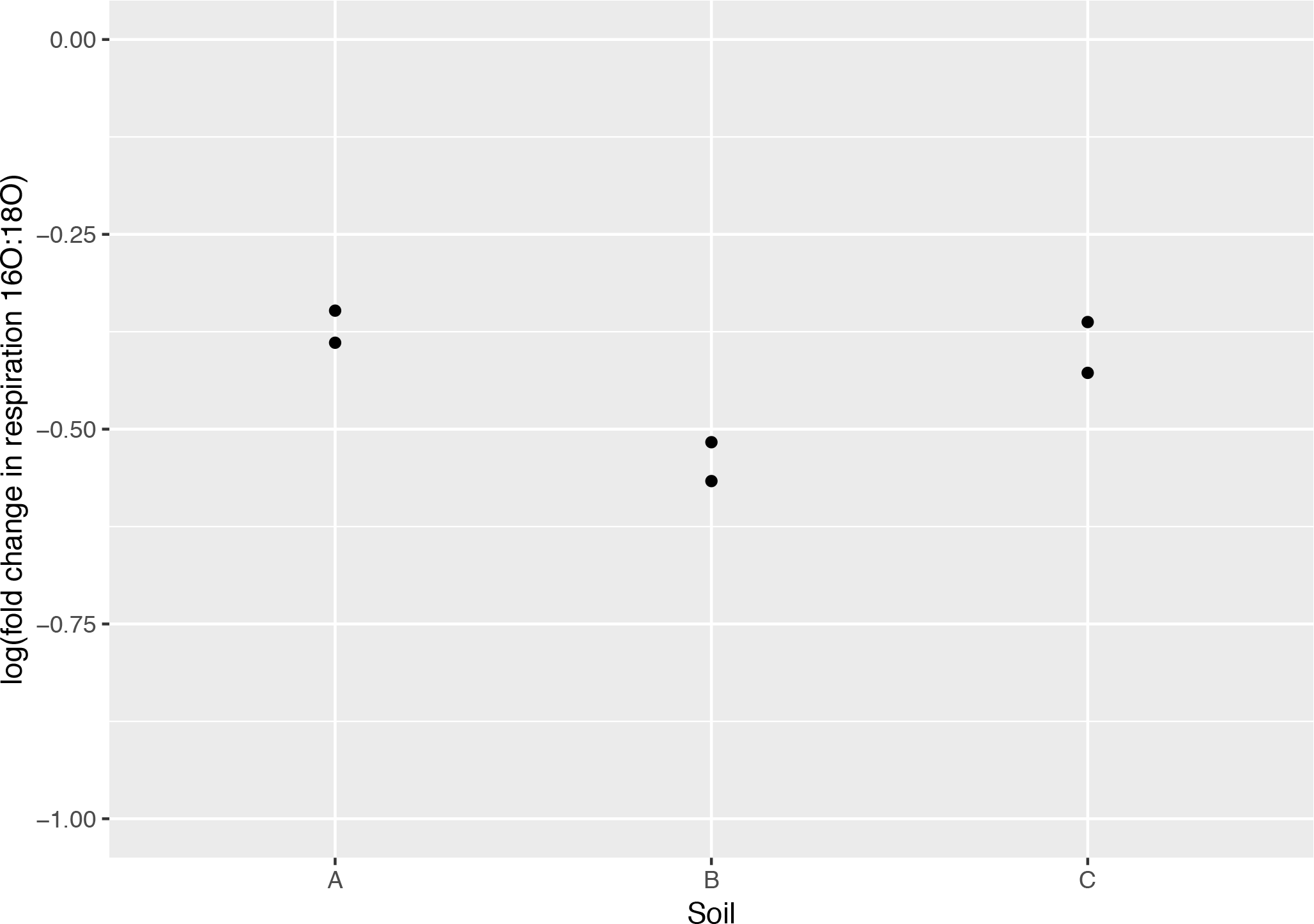
Respiration suppression after addition of 96 at% ^18^O-water compared to ^16^O-water.

## References

[1] Steven D Allison, Matthew D Wallenstein, and Mark A Bradford. Soil-carbon response to warming dependent on microbial physiology. Nature Geoscience, 3(5):336, 2010.

[2] Jianwei Li, Gangsheng Wang, Steven D Allison, Melanie A Mayes, and Yiqi Luo. Soil carbon sensitivity to temperature and carbon use efficiency compared across microbial-ecosystem models of varying complexity. Biogeochemistry, 119(1-3):67–84, 2014.

[3] Serita D Frey, Juhwan Lee, Jerry M Melillo, and Johan Six. The temperature response of soil microbial efficiency and its feedback to climate. Nature Climate Change, 3(4):395, 2013.

[4] Seeta A Sistla, Edward B Rastetter, and Joshua P Schimel. Responses of a tundra system to warming using scamps: a stoichiometrically coupled, acclimating microbe–plant–soil model. Ecological Monographs, 84(1):151–170, 2014.

[5] Kevin M Geyer, Paul Dijkstra, Robert Sinsabaugh, and Serita D Frey. Clarifying the interpretation of carbon use efficiency in soil through methods comparison. Soil Biology and Biochemistry, 128:79–88, 2019.

[6] Shannon B Hagerty, Steven D Allison, and Joshua P Schimel. Evaluating soil microbial carbon use efficiency explicitly as a function of cellular processes: implications for measurements and models. Biogeochemistry, 140(3):269–283, 2018.

[7] PJF Gommers, BJ Van Schie, JP Van Dijken, and JG Kuenen. Biochemical limits to microbial growth yields: an analysis of mixed substrate utilization. Biotechnology and bioengineering, 32(1):86–94, 1988.

[8] CA Lehmeier, F Ballantyne IV, K Min, and SA Billings. Temperature-mediated changes in microbial carbon use efficiency and 13 c discrimination. Biogeosciences Discussions, 12(20), 2015.

[9] Marie Spohn, Erich M Pötsch, Stephanie A Eichorst, Dagmar Woebken, Wolfgang Wanek, and Andreas Richter. Soil microbial carbon use efficiency and biomass turnover in a long-term fertilization experiment in a temperate grassland. Soil Biology and Biochemistry, 97:168–175, 2016.

[10] Christopher Poeplau, Mirjam Helfrich, Rene Dechow, Márton Szoboszlay, Christoph C Tebbe, Axel Don, Bärbel Greiner, Dorit Zopf, Ulrich Thumm, Hein Korevaar, et al. Increased microbial anabolism contributes to soil carbon sequestration by mineral fertilization in temperate grasslands. Soil Biology and Biochemistry, 130:167–176, 2019.

[11] Tom WN Walker, Christina Kaiser, Florian Strasser, Craig W Herbold, Niki IW Leblans, Dagmar Woebken, Ivan A Janssens, Bjarni D Sigurdsson, and Andreas Richter. Microbial temperature sensitivity and biomass change explain soil carbon loss with warming. Nature climate change, 8(10):885, 2018.

[12] Marie Spohn, Karoline Klaus, Wolfgang Wanek, and Andreas Richter. Microbial carbon use efficiency and biomass turnover times depending on soil depth–implications for carbon cycling. Soil Biology and Biochemistry, 96:74–81, 2016.

[13] R Core Team. R: A Language and Environment for Statistical Computing. R Foundation for Statistical Computing, Vienna, Austria, 2016.

[14] Hadley Wickham. ggplot2: Elegant Graphics for Data Analysis. Springer-Verlag New York, 2009.

[15] Hadley Wickham. The split-apply-combine strategy for data analysis. Journal of Statistical Software, 40(1):1–29, 2011.

[16] Winston Chang, Joe Cheng, JJ Allaire, Yihui Xie, and Jonathan McPherson. shiny: Web Application Framework for R, 2018. R package version 1.1.0.

[17] Alboukadel Kassambara. ggpubr: ‘ggplot2’ Based Publication Ready Plots, 2018. R package version 0.2.

[18] Bradley S. Stevenson, Stephanie A. Eichorst, John T. Wertz, Thomas M. Schmidt, and John A. Breznak. New strategies for cultivation and detection of previously uncultured microbes. Applied and Environmental Microbiology, 70(8):4748–4755, 2004.

[19] E C. Uitterlinden A G. Muyzer, G. de Waal. Profiling of complex microbial populations by denaturing gradient gel electrophoresis analysis of polymerase chain reaction-amplified genes coding for 16s rrna.

[20] Noah Fierer, Jason A Jackson, Rytas Vilgalys, and Robert B Jackson. Assessment of soil microbial community structure by use of taxon-specific quantitative pcr assays. Appl. Environ. Microbiol., 71(7):4117–4120, 2005.

[21] Lotus A Lofgren, Jessie K Uehling, Sara Branco, Thomas D Bruns, Francis Martin, and Peter G Kennedy. Genome-based estimates of fungal rdna copy number variation across phylogenetic scales and ecological lifestyles. Molecular ecology, 2018.

[22] Kristen M DeAngelis, Grace Pold, Begöm D Topçuoğlu, Linda TA van Diepen, Rebecca M Varney, Jeffrey L Blanchard, Jerry Melillo, and Serita D Frey. Long-term forest soil warming alters microbial communities in temperate forest soils. Frontiers in microbiology, 6:104, 2015.

[23] Katerina Papp, Rebecca L Mau, Michaela Hayer, Benjamin J Koch, Bruce A Hungate, and Egbert Schwartz. Quantitative stable isotope probing with h2 18 o reveals that most bacterial taxa in soil synthesize new ribosomal rna. The ISME journal, 12(12):3043–3045, 2018.

[24] Jay T Lennon and Stuart E Jones. Microbial seed banks: the ecological and evolutionary implications of dormancy. Nature reviews microbiology, 9(2):119, 2011.

[25] Gangsheng Wang, Melanie A Mayes, Lianhong Gu, and Christopher W Schadt. Representation of dormant and active microbial dynamics for ecosystem modeling. PloS one, 9(2):e89252, 2014.

[26] Helen W Kreuzer-Martin, James R Ehleringer, and Eric L Hegg. Oxygen isotopes indicate most intracellular water in log-phase escherichia coli is derived from metabolism. Proceedings of the National Academy of Sciences, 102(48):17337–17341, 2005.

[27] Helen W Kreuzer-Martin, Michael J Lott, James R Ehleringer, and Eric L Hegg. Metabolic processes account for the majority of the intracellular water in log-phase escherichia coli cells as revealed by hydrogen isotopes. Biochemistry, 45(45):13622–13630, 2006.

[28] Hui Li, Chan Yu, Fei Wang, Sae Jung Chang, Jun Yao, and Ruth E Blake. Probing the metabolic water contribution to intracellular water using oxygen isotope ratios of po4. Proceedings of the National Academy of Sciences, 113(21):5862–5867, 2016.

[29] Bruce A Hungate, Rebecca L Mau, Egbert Schwartz, J Gregory Caporaso, Paul Dijkstra, Natasja van Gestel, Benjamin J Koch, Cindy M Liu, Theresa A McHugh, Jane C Marks, et al. Quantitative microbial ecology through stable isotope probing. Applied and environmental microbiology, pages AEM–02280, 2015.

[30] Zachary T Aanderud and Jay T Lennon. Validation of heavy-water stable isotope probing for the characterization of rapidly responding soil bacteria. Applied and environmental microbiology, pages AEM–02735, 2011.

[31] Larry M Feinstein, Woo Jun Sul, and Christopher B Blackwood. Assessment of bias associated with incomplete extraction of microbial dna from soil. Appl. Environ. Microbiol., 75(16):5428–5433, 2009.

[32] Steven J Blazewicz, Egbert Schwartz, and Mary K Firestone. Growth and death of bacteria and fungi underlie rainfall-induced carbon dioxide pulses from seasonally dried soil. Ecology, 95(5):1162–1172, 2014.

[33] Johannes Rousk and Erland Bååth. Growth of saprotrophic fungi and bacteria in soil. FEMS Microbiology Ecology, 78(1):17–30, 2011.

[34] Benjamin J Koch, Theresa A McHugh, Michaela Hayer, Egbert Schwartz, Steven J Blazewicz, Paul Dijkstra, Natasja Gestel, Jane C Marks, Rebecca L Mau, Ember M Morrissey, et al. Estimating taxon-specific population dynamics in diverse microbial communities. Ecosphere, 9(1), 2018.

[35] Margarida Soares and Johannes Rousk. Microbial growth and carbon use efficiency in soil: Links to fungal-bacterial dominance, soc-quality and stoichiometry. Soil Biology and Biochemistry, 2019.

[36] Beth Gibson, Daniel J Wilson, Edward Feil, and Adam Eyre-Walker. The distribution of bacterial doubling times in the wild. Proceedings of the Royal Society B: Biological Sciences, 285(1880):20180789, 2018.

[37] Barbara Drigo, Ian C Anderson, GSK Kannangara, John WG Cairney, and David Johnson. Rapid incorporation of carbon from ectomycorrhizal mycelial necromass into soil fungal communities. Soil Biology and Biochemistry, 49:4–10, 2012.

[38] Lars Reier Bakken and Rolf Arnt Olsen. Dna-content of soil bacteria of different cell size. Soil Biology and Biochemistry, 21(6):789–793, 1989.

[39] Céline Mouginot, Rika Kawamura, Kristin L Matulich, Renaud Berlemont, Steven D Allison, Anthony S Amend, and Adam C Martiny. Elemental stoichiometry of fungi and bacteria strains from grassland leaf litter. Soil Biology and Biochemistry, 76:278–285, 2014.

[40] Maria C Portillo, Jonathan W Leff, Christian L Lauber, and Noah Fierer. Cell size distributions of soil bacterial and archaeal taxa. Appl. Environ. Microbiol., 79(24):7610–7617, 2013.

[41] Anja Kristiansen, Aaron M Saunders, Aviaja A Hansen, Per H Nielsen, and Jeppe L Nielsen. Community structure of bacteria and fungi in aerosols of a pig confinement building. FEMS microbiology ecology, 80(2):390–401, 2012.

[42] H Christensen, RA Olsen, and LR Bakken. Flow cytometric measurements of cell volumes and dna contents during culture of indigenous soil bacteria. Microbial ecology, 29(1):49–62, 1995.

[43] Henrik Christensen, Lars R Bakken, and Rolf A Olsen. Soil bacterial dna and biovolume profiles measured by flow-cytometry. FEMS Microbiology Ecology, 11(3-4):129–140, 1993.

[44] W Makino, JB Cotner, RW Sterner, and JJ Elser. Are bacteria more like plants or animals? growth rate and resource dependence of bacterial c: N: P stoichiometry. Functional Ecology, 17(1):121–130, 2003.

[45] Traute-Heidi Anderson and Rainer Martens. Dna determinations during growth of soil microbial biomasses. Soil Biology and Biochemistry, 57:487–495, 2013.

[46] IJ Grimmett, KN Shipp, A Macneil, and F Bärlocher. Does the growth rate hypothesis apply to aquatic hyphomycetes? fungal ecology, 6(6):493–500, 2013.

[47] Sara E Leckie, Cindy E Prescott, Susan J Grayston, Josh D Neufeld, and William W Mohn. Comparison of chloroform fumigation-extraction, phospholipid fatty acid, and dna methods to determine microbial biomass in forest humus. Soil Biology and Biochemistry, 36(3):529–532, 2004.

[48] DS Jenkinson. The effects of biocidal treatments on metabolism in soil—iv. the decomposition of fumigated organisms in soil. Soil Biology and Biochemistry, 8(3):203–208, 1976.

[49] KR Tate, DJ Ross, and CW Feltham. A direct extraction method to estimate soil microbial c: effects of experimental variables and some different calibration procedures. Soil Biology and Biochemistry, 20(3):329–335, 1988.

[50] Marie-Christine Dictor, Laurent Tessier, and Guy Soulas. Reassessement of the kec coeffcient of the fumigation–extraction method in a soil profile. Soil Biology and Biochemistry, 30(2):119–127, 1998.

[51] Rainer Georg Joergensen. The fumigation-extraction method to estimate soil microbial biomass: calibration of the kec value. Soil Biology and Biochemistry, 28(1):25–31, 1996.

[52] JP Martin, JO Ervin, and RA Shepherd. Decomposition and aggregating effect of fungus cell material in soil 1. Soil Science Society of America Journal, 23(3):217–220, 1959.

[53] Steven F Stoddard, Byron J Smith, Robert Hein, Benjamin RK Roller, and Thomas M Schmidt. rrn db: improved tools for interpreting rrna gene abundance in bacteria and archaea and a new foundation for future development. Nucleic acids research, 43(D1):D593–D598, 2014.

[54] SM Dineen, Rt Aranda, DL Anders, and JM Robertson. An evaluation of commercial dna extraction kits for the isolation of bacterial spore dna from soil. Journal of applied microbiology, 109(6):1886–1896, 2010.

[55] JT Lennon, ME Muscarella, SA Placella, and BK Lehmkuhl. How, when, and where relic dna affects microbial diversity. mBio, 9(3):e00637–18, 2018.

[56] XIAO Hai-Feng, LI Gen, LI Da-Ming, HU Feng, and LI Hui-Xin. Effect of different bacterial-feeding nematode species on soil bacterial numbers, activity, and community composition. Pedosphere, 24(1):116–124, 2014.

[57] Cyril Blanc, M Sy, D Djigal, Alain Brauman, P Normand, and Cécile Villenave. Nutrition on bacteria by bacterial-feeding nematodes and consequences on the structure of soil bacterial community. European Journal of Soil Biology, 42:S70–S78, 2006.

[58] I-Min A Chen, Victor M Markowitz, Ken Chu, Krishna Palaniappan, Ernest Szeto, Manoj Pillay, Anna Ratner, Jinghua Huang, Evan Andersen, Marcel Huntemann, et al. Img/m: integrated genome and metagenome comparative data analysis system. Nucleic acids research, page gkw929, 2016.

[59] Daniel S Alessi, Dana M Walsh, and Jeremy B Fein. Uncertainties in determining microbial biomass c using the chloroform fumigation–extraction method. Chemical Geology, 280(1-2):58–64, 2011.

[60] Raj Setia, Suman Lata Verma, and Petra Marschner. Measuring microbial biomass carbon by direct extraction–comparison with chloroform fumigation-extraction. European journal of soil biology, 53:103–106, 2012.

[61] Ruth E Blake, Aleksandr V Surkov, Lisa M Stout, Hui Li, Sae Jung Chang, Deb P Jaisi, Albert S Colman, and Yuhong Liang. Dna thermometry: A universal biothermometer in the 18o/16o ratio of po4 in dna. American Journal of Science, 316(9):813–838, 2016.

[62] Grace Pold, A Stuart Grandy, Jerry M Melillo, and Kristen M DeAngelis. Changes in substrate availability drive carbon cycle response to chronic warming. Soil Biology and Biochemistry, 110:68–78, 2017.

[63] Noah Fierer, Michael S Strickland, Daniel Liptzin, Mark A Bradford, and Cory C Cleveland. Global patterns in belowground communities. Ecology letters, 12(11):1238–1249, 2009.

[64] Petr Baldrian, Miroslav Kolařík, Martina Štursová, Jan Kopeckỳ, Vendula Valášková, Tomáš Vĕtrovskỳ, Lucia Žifčáková, Jaroslav Šnajdr, Jakub Rídl, Čestmír Vlček, et al. Active and total microbial communities in forest soil are largely different and highly stratified during decomposition. The ISME journal, 6(2):248, 2012.

[65] Alex Silva-Sánchez, Margarida Soares, and Johannes Rousk. Testing the dependence of microbial growth and carbon use efficiency on nitrogen availability, ph, and organic matter quality. Soil Biology and Biochemistry, 2019.

[66] Mark A Lever, Karyn L Rogers, Karen G Lloyd, Jörg Overmann, Bernhard Schink, Rudolf K Thauer, Tori M Hoehler, and Bo Barker Jørgensen. Life under extreme energy limitation: a synthesis of laboratory-and field-based investigations. FEMS microbiology reviews, 39(5):688–728, 2015.

[67] William D Donachie. Relationship between cell size and time of initiation of dna replication. Nature, 219(5158):1077, 1968.

[68] HE Kubitschek. Constancy of the ratio of dna to cell volume in steady-state cultures of escherichia coli br. Biophysical journal, 14(2):119, 1974.

