## supplementary file S3 - R script for "Heavy and wet: evaluating the validity and implications of assumptions made when measuring growth efficiency using ^18^O water"

#sensitivity analysis of CUE calcs using mean and extreme values observed in the literature:

outy<-matrix(ncol=16, nrow=100000)
i<-1
for(FBratio in c(0.1, 0.001, 0.999)){
  for(MBCDNAratB in c(13.22,3.6,37)){
    for(MBCDNAratF in c(1070, 90, 3300)){
      for(DNAexteffF in c(0.33, 0.04, 0.5)){
        for(DNAexteffB in c(0.5, 0.06, 0.5)){
          for(MBCexteffF in c(0.33, 0.21, 0.45)){
            for(MBCexteffB in c( 0.32, 0.10, 0.51)){
              for(DNAbias in c(0.7, 0.3, 1)){
                for(GRbact in c(0.167, 0.003, 3.4)){
                  for(GRfun in c(0.0088, 0.0001, 0.8)){
                    
  totalDNA<-30*FBratio+3
  #actual MBC:DNA ratio
  MBC<-FBratio*totalDNA*as.numeric(MBCDNAratF)+(1-FBratio)*totalDNA*as.numeric(MBCDNAratB)
  MBCDNA<-MBC/totalDNA
  #observed MBC:DNA ratio if only DNA extraction bias
  totalDNAd<-FBratio*totalDNA*as.numeric(DNAexteffF)+(1-FBratio)*totalDNA*as.numeric(DNAexteffB)
  MBCDNAd<-MBC/totalDNAd
  #Observed MBC:DNA ratio if only MBC extraction bias
  MBCc<-FBratio*totalDNA*as.numeric(MBCDNAratF)*as.numeric(MBCexteffF)+(1-FBratio)*totalDNA*as.numeric(MBCDNAratB)*as.numeric(MBCexteffB)
  MBCDNAc<-MBCc/totalDNA
  #Observed MBC:DNA ratio if both DNA and MBC extraction biases
  MBCDNAdc<-MBCc/totalDNAd
  
  #True MBC growth if both bacteria and fungi grow at community level mean
  GRmean<-FBratio*as.numeric(GRfun)+as.numeric(GRbact)*(1-FBratio)
  
  True18Osame<-GRmean*totalDNA
  TrueMBCsame<-GRmean*totalDNA*FBratio*as.numeric(MBCDNAratF)+GRmean*totalDNA*(1-FBratio)*as.numeric(MBCDNAratB)
  #observed MBC growth if both bacteria and fungi grow at 0.0104 day-1, under various extraction biases
  outy[i,11]<-MBCsamed<-(as.numeric(DNAbias)*(GRmean*totalDNA*FBratio*as.numeric(DNAexteffF)*MBCDNAd+GRmean*totalDNA*(1-FBratio)*as.numeric(DNAexteffB)*MBCDNAd))/TrueMBCsame
  outy[i,12]<-MBCsamec<-(as.numeric(DNAbias)*(GRmean*totalDNA*FBratio*MBCDNAc+GRmean*totalDNA*(1-FBratio)*MBCDNAc))/TrueMBCsame
  outy[i,13]<-MBCsamedc<-(as.numeric(DNAbias)*(GRmean*totalDNA*FBratio*as.numeric(DNAexteffF)*MBCDNAdc+GRmean*totalDNA*(1-FBratio)*as.numeric(DNAexteffB)*MBCDNAdc))/TrueMBCsame
  
  #True MBC growth if bacteria grow at 0.12 day-1 and fungi at 0.0088 day -1
  
  TrueMBCdiff<-(as.numeric(GRfun)*totalDNA*FBratio*as.numeric(MBCDNAratF)+as.numeric(GRbact)*totalDNA*(1-FBratio)*as.numeric(MBCDNAratB))
  #observed MBC growth if bacteria grow at 0.12 day-1 and fungi at 0.0088 day -1, under various extraction biases
  outy[i,14]<-MBCdiffd<-(as.numeric(DNAbias)*(as.numeric(GRfun)*totalDNA*FBratio*as.numeric(DNAexteffF)*MBCDNAd+as.numeric(GRbact)*totalDNA*(1-FBratio)*as.numeric(DNAexteffB)*MBCDNAd))/TrueMBCdiff
  outy[i,15]<-MBCdiffc<-(as.numeric(DNAbias)*(as.numeric(GRfun)*totalDNA*FBratio*MBCDNAc+as.numeric(GRbact)*totalDNA*(1-FBratio)*MBCDNAc))/TrueMBCdiff
  outy[i,16]<-MBCdiffdc<-(as.numeric(DNAbias)*(as.numeric(GRfun)*totalDNA*FBratio*as.numeric(DNAexteffF)*MBCDNAdc+as.numeric(GRbact)*totalDNA*(1-FBratio)*as.numeric(DNAexteffB)*MBCDNAdc))/TrueMBCdiff
  
  outy[i,1]<-MBCDNAratB
    outy[i,2]<-MBCDNAratF
    outy[i,3]<-DNAexteffF
      outy[i,4]<-DNAexteffB
    outy[i,5]<-MBCexteffF
    outy[i,6]<-MBCexteffB
    outy[i,7]<-DNAbias
    outy[i,8]<-GRbact
    outy[i,9]<-GRfun
    outy[i,10]<-FBratio
    i<-i+1
                  } }}}}}}}}}


outy<-as.data.frame(outy)
outy<-subset(outy, !is.na(outy$V1))
names(outy)<-c("MBCDNAratB", "MBCDNAratF", "DNAexteffF","DNAexteffB","MBCexteffF","MBCexteffB","DNAbias", "GRbact", "GRfun", "FBratio", "MBCsamed", "MBCsamec", "MBCsamedc",  "MBCdiffd","MBCdiffc","MBCdiffdc")

reference <- c( 13.22,1070,0.33,0.5, 0.33,0.32,0.7,0.167,0.0088,0.1)#defining parameters to keep for all runs
outysubs<- outy[apply(outy[,1:10], 1, function(x) sum(reference  == x) >= 9), ]   #subsetting so only get rows where a single value is changed           
outylog<-log10(outysubs)#taking logarithm of all the values in the table
outylog2<-unique( outylog )

outy0.001FB<-subset(outy, FBratio==0.001)
outy0.999FB<-subset(outy, FBratio==0.999)
reference2 <- c( 13.22,1070,0.33,0.5, 0.33,0.32,0.7,0.167,0.0088)#defining parameters to keep for all runs
outysubs0.001FB<- outy0.001FB[apply(outy0.001FB[,1:9], 1, function(x) sum(reference2  == x) >= 8), ]   #subsetting so only get rows where a single value is changed           
outysubs0.999FB<- outy0.999FB[apply(outy0.999FB[,1:9], 1, function(x) sum(reference2  == x) >= 8), ]   #subsetting so only get rows where a single value is changed           

outylog0.001FB<-log10(outysubs0.001FB)#taking logarithm of all the values in the table
outylog0.001FB2<-unique( outylog0.001FB )
outylog0.999FB<-log10(outysubs0.999FB)#taking logarithm of all the values in the table
outylog0.999FB2<-unique( outylog0.999FB )

write.table(outylog0.999FB2, "~/Desktop/LuizMicrocosms/CUEsensitivityAnalysis190207log100.999FB.txt", sep="\t",row.names = FALSE)
write.table(outylog0.001FB2, "~/Desktop/LuizMicrocosms/CUEsensitivityAnalysis190207log100.001FB.txt", sep="\t",row.names = FALSE)


#write.table(outylog2, "~/Desktop/LuizMicrocosms/CUEsensitivityAnalysis190207log10.txt", sep="\t",row.names = FALSE)


abs(log10(13.22))#MBCDNAratB
  abs(log10(1070))#MBCDNAratF
    abs(log10(0.33))#DNAexteffF 
    abs(log10(0.5))#DNAexteffB
    abs(log10(0.33))#MBCexteffF
          abs(log10(0.32))#MBCexteffB
            abs(log10(1))#DNAbias
              abs(log10(0.167))#GRbact
                abs(log10(0.0088))#GRfun
                abs(log10(0.1))#FBratio   
                
 
sensival<-read.delim("~/Desktop/LuizMicrocosms/CUEsensitivityAnalysis190207summary.txt")
sensival$MBCmeasure<-factor(sensival$MBCmeasure, levels=c("MBCsamed","MBCsamec","MBCsamedc", "MBCdiffd","MBCdiffc", "MBCdiffdc"))
sensival$MBCmeasure<-gsub(pattern = "MBC", "", sensival$MBCmeasure)
ggplot(subset(sensival, Fbrat==0.1), aes(x=parameter, y=sensitivity, colour=parameter))+geom_point(position="dodge")+theme_bw()+facet_wrap(~MBCmeasure)+ theme(panel.border = element_blank(), panel.grid.major = element_blank(),panel.grid.minor = element_blank(), axis.line = element_line(colour = "black"))                   
ggplot(subset(sensival, Fbrat==0.1), aes(x=MBCmeasure, y=sensitivity, colour=parameter, shape=parameter))+geom_point(position="jitter", size=5)+theme_bw()+ theme(panel.border = element_blank(), panel.grid.major = element_blank(),panel.grid.minor = element_blank(), axis.line = element_line(colour = "black"),axis.text=element_text(size=12),axis.title=element_text(size=12))+
  ylab("sensitivity value")+xlab("extraction bias scenario")+scale_shape_manual(values=c(0:10))
ggplot(subset(sensival, MBCmeasure=="diffdc"&!parameter=="FBratio"), aes(x=Fbrat, y=sensitivity, colour=parameter))+geom_point()+theme_bw()+ theme(panel.border = element_blank(), panel.grid.major = element_blank(),panel.grid.minor = element_blank(), axis.line = element_line(colour = "black"),axis.text=element_text(size=11),axis.title=element_text(size=11))+
  ylab("sensitivity value")+xlab("fungal:bacterial DNA ratio")+scale_shape_manual(values=c(0:10))+facet_wrap(~parameter)+scale_x_continuous(breaks = seq(0, 1, by = 0.5))
ggplot(subset(sensival, MBCmeasure=="diffdc"&!parameter=="FBratio"), aes(x=Fbrat, y=sensitivity, colour=parameter))+geom_smooth()+theme_bw()+ theme(panel.border = element_blank(), panel.grid.major = element_blank(),panel.grid.minor = element_blank(), axis.line = element_line(colour = "black"),axis.text=element_text(size=11),axis.title=element_text(size=11))+
  ylab("sensitivity value")+xlab("fungal:bacterial DNA ratio")+scale_shape_manual(values=c(0:10))+facet_wrap(~parameter)+scale_x_continuous(breaks = seq(0, 1, by = 0.5))

gg_color_hue <- function(n) {
  hues = seq(15, 375, length = n + 1)
  hcl(h = hues, l = 65, c = 100)[1:n]
}#redefning colours so can match parameters in both figures
cols = gg_color_hue(10)[c(1:3,5:10)]
ggplot(subset(sensival, MBCmeasure=="diffdc"&!parameter=="FBratio"), aes(x=Fbrat, y=sensitivity, colour=parameter))+geom_point(size=5)+theme_bw()+ theme(panel.border = element_blank(), panel.grid.major = element_blank(),panel.grid.minor = element_blank(), axis.line = element_line(colour = "black"),axis.text=element_text(size=14),axis.title=element_text(size=14))+
  ylab("sensitivity value")+xlab("fungal:bacterial DNA ratio")+scale_shape_manual(values=c(0:10))+facet_wrap(~parameter)+scale_x_continuous(breaks = seq(0, 1, by = 0.5))

ggplot(subset(sensival, MBCmeasure=="diffdc"&!parameter=="FBratio"), aes(x=Fbrat, y=sensitivity, colour=parameter))+geom_line()+theme_bw()+ theme(panel.border = element_blank(), panel.grid.major = element_blank(),panel.grid.minor = element_blank(), axis.line = element_line(colour = "black"),axis.text=element_text(size=11),axis.title=element_text(size=11))+
  ylab("sensitivity value")+xlab("fungal:bacterial DNA ratio")+scale_shape_manual(values=c(0:10))+scale_x_continuous(breaks = seq(0, 1, by = 0.5))
ggplot(subset(sensival, MBCmeasure=="diffdc"&!parameter=="FBratio"), aes(x=Fbrat, y=sensitivity, colour=parameter))+geom_point()+theme_bw()+ theme(panel.border = element_blank(), panel.grid.major = element_blank(),panel.grid.minor = element_blank(), axis.line = element_line(colour = "black"),axis.text=element_text(size=11),axis.title=element_text(size=11))+
  ylab("sensitivity value")+xlab("fungal:bacterial DNA ratio")+scale_shape_manual(values=c(0:10))+scale_x_continuous(breaks = seq(0, 1, by = 0.5))


#REPEAT, only with the literature value halved or doubled

outy<-matrix(ncol=16, nrow=100000)
i<-1
for(FBratio in c(0.1, 0.05, 0.2)){
  for(MBCDNAratB in c(13.22,6.61,26.44)){
    for(MBCDNAratF in c(1070, 535, 2140)){
      for(DNAexteffF in c(0.33, 0.165, 0.66)){
        for(DNAexteffB in c(0.5, 0.25, 1)){
          for(MBCexteffF in c(0.33, 0.165, 0.66)){
            for(MBCexteffB in c( 0.32, 0.16, 0.64)){
              for(DNAbias in c(0.7)){
                for(GRbact in c(0.167, .0835, .334)){
                  for(GRfun in c(0.0088, 0.0044, 0.0176)){
                    
                    totalDNA<-30*FBratio+3
                    #actual MBC:DNA ratio
                    MBC<-FBratio*totalDNA*as.numeric(MBCDNAratF)+(1-FBratio)*totalDNA*as.numeric(MBCDNAratB)
                    MBCDNA<-MBC/totalDNA
                    #observed MBC:DNA ratio if only DNA extraction bias
                    totalDNAd<-FBratio*totalDNA*as.numeric(DNAexteffF)+(1-FBratio)*totalDNA*as.numeric(DNAexteffB)
                    MBCDNAd<-MBC/totalDNAd
                    #Observed MBC:DNA ratio if only MBC extraction bias
                    MBCc<-FBratio*totalDNA*as.numeric(MBCDNAratF)*as.numeric(MBCexteffF)+(1-FBratio)*totalDNA*as.numeric(MBCDNAratB)*as.numeric(MBCexteffB)
                    MBCDNAc<-MBCc/totalDNA
                    #Observed MBC:DNA ratio if both DNA and MBC extraction biases
                    MBCDNAdc<-MBCc/totalDNAd
                    
                    #True MBC growth if both bacteria and fungi grow at community level mean
                    GRmean<-FBratio*as.numeric(GRfun)+as.numeric(GRbact)*(1-FBratio)
                    
                    True18Osame<-GRmean*totalDNA
                    TrueMBCsame<-GRmean*totalDNA*FBratio*as.numeric(MBCDNAratF)+GRmean*totalDNA*(1-FBratio)*as.numeric(MBCDNAratB)
                    #observed MBC growth if both bacteria and fungi grow at 0.0104 day-1, under various extraction biases
                    outy[i,11]<-MBCsamed<-(as.numeric(DNAbias)*(GRmean*totalDNA*FBratio*as.numeric(DNAexteffF)*MBCDNAd+GRmean*totalDNA*(1-FBratio)*as.numeric(DNAexteffB)*MBCDNAd))/TrueMBCsame
                    outy[i,12]<-MBCsamec<-(as.numeric(DNAbias)*(GRmean*totalDNA*FBratio*MBCDNAc+GRmean*totalDNA*(1-FBratio)*MBCDNAc))/TrueMBCsame
                    outy[i,13]<-MBCsamedc<-(as.numeric(DNAbias)*(GRmean*totalDNA*FBratio*as.numeric(DNAexteffF)*MBCDNAdc+GRmean*totalDNA*(1-FBratio)*as.numeric(DNAexteffB)*MBCDNAdc))/TrueMBCsame
                    
                    #True MBC growth if bacteria grow at 0.12 day-1 and fungi at 0.0088 day -1
                    
                    TrueMBCdiff<-(as.numeric(GRfun)*totalDNA*FBratio*as.numeric(MBCDNAratF)+as.numeric(GRbact)*totalDNA*(1-FBratio)*as.numeric(MBCDNAratB))
                    #observed MBC growth if bacteria grow at 0.12 day-1 and fungi at 0.0088 day -1, under various extraction biases
                    outy[i,14]<-MBCdiffd<-(as.numeric(DNAbias)*(as.numeric(GRfun)*totalDNA*FBratio*as.numeric(DNAexteffF)*MBCDNAd+as.numeric(GRbact)*totalDNA*(1-FBratio)*as.numeric(DNAexteffB)*MBCDNAd))/TrueMBCdiff
                    outy[i,15]<-MBCdiffc<-(as.numeric(DNAbias)*(as.numeric(GRfun)*totalDNA*FBratio*MBCDNAc+as.numeric(GRbact)*totalDNA*(1-FBratio)*MBCDNAc))/TrueMBCdiff
                    outy[i,16]<-MBCdiffdc<-(as.numeric(DNAbias)*(as.numeric(GRfun)*totalDNA*FBratio*as.numeric(DNAexteffF)*MBCDNAdc+as.numeric(GRbact)*totalDNA*(1-FBratio)*as.numeric(DNAexteffB)*MBCDNAdc))/TrueMBCdiff
                    
                    outy[i,1]<-MBCDNAratB
                    outy[i,2]<-MBCDNAratF
                    outy[i,3]<-DNAexteffF
                    outy[i,4]<-DNAexteffB
                    outy[i,5]<-MBCexteffF
                    outy[i,6]<-MBCexteffB
                    outy[i,7]<-DNAbias
                    outy[i,8]<-GRbact
                    outy[i,9]<-GRfun
                    outy[i,10]<-FBratio
                    i<-i+1
                  } }}}}}}}}}


outy<-as.data.frame(outy)
outy<-subset(outy, !is.na(outy$V1))
names(outy)<-c("MBCDNAratB", "MBCDNAratF", "DNAexteffF","DNAexteffB","MBCexteffF","MBCexteffB","DNAbias", "GRbact", "GRfun", "FBratio", "MBCsamed", "MBCsamec", "MBCsamedc",  "MBCdiffd","MBCdiffc","MBCdiffdc")

reference <- c( 13.22,1070,0.33,0.5, 0.33,0.32,0.7,0.167,0.0088,0.1)#defining parameters to keep for all runs
outysubs<- outy[apply(outy[,1:10], 1, function(x) sum(reference  == x) >= 9), ]   #subsetting so only get rows where a single value is changed           

outylog<-log10(outysubs)#taking logarithm of all the values in the table
outylog2<-unique( outylog )
write.table(outylog2, "~/Desktop/LuizMicrocosms/CUEsensitivityAnalysis190207log10_halve.txt", sep="\t",row.names = FALSE)


###########
outy<-matrix(ncol=16, nrow=160000)
i<-1
for(FBratio in c(0.2,0.3,0.4,0.5,0.6,0.7,0.8,0.9)){
  for(FBratio in c(0.01,0.05,0.95,0.99,0.25)){
  for(MBCDNAratB in c(13.22,3.6,37)){
    for(MBCDNAratF in c(1070, 90, 3300)){
      for(DNAexteffF in c(0.33, 0.04, 0.5)){
        for(DNAexteffB in c(0.5, 0.06, 0.5)){
          for(MBCexteffF in c(0.33, 0.21, 0.45)){
            for(MBCexteffB in c( 0.32, 0.10, 0.51)){
              for(DNAbias in c(0.7, 0.3, 1)){
                for(GRbact in c(0.167, 0.003, 3.4)){
                  for(GRfun in c(0.0088, 0.0001, 0.8)){
                    
                    totalDNA<-30*FBratio+3
                    #actual MBC:DNA ratio
                    MBC<-FBratio*totalDNA*as.numeric(MBCDNAratF)+(1-FBratio)*totalDNA*as.numeric(MBCDNAratB)
                    MBCDNA<-MBC/totalDNA
                    #observed MBC:DNA ratio if only DNA extraction bias
                    totalDNAd<-FBratio*totalDNA*as.numeric(DNAexteffF)+(1-FBratio)*totalDNA*as.numeric(DNAexteffB)
                    MBCDNAd<-MBC/totalDNAd
                    #Observed MBC:DNA ratio if only MBC extraction bias
                    MBCc<-FBratio*totalDNA*as.numeric(MBCDNAratF)*as.numeric(MBCexteffF)+(1-FBratio)*totalDNA*as.numeric(MBCDNAratB)*as.numeric(MBCexteffB)
                    MBCDNAc<-MBCc/totalDNA
                    #Observed MBC:DNA ratio if both DNA and MBC extraction biases
                    MBCDNAdc<-MBCc/totalDNAd
                    
                    #True MBC growth if both bacteria and fungi grow at community level mean
                    GRmean<-FBratio*as.numeric(GRfun)+as.numeric(GRbact)*(1-FBratio)
                    
                    True18Osame<-GRmean*totalDNA
                    TrueMBCsame<-GRmean*totalDNA*FBratio*as.numeric(MBCDNAratF)+GRmean*totalDNA*(1-FBratio)*as.numeric(MBCDNAratB)
                    #observed MBC growth if both bacteria and fungi grow at 0.0104 day-1, under various extraction biases
                    outy[i,11]<-MBCsamed<-(as.numeric(DNAbias)*(GRmean*totalDNA*FBratio*as.numeric(DNAexteffF)*MBCDNAd+GRmean*totalDNA*(1-FBratio)*as.numeric(DNAexteffB)*MBCDNAd))/TrueMBCsame
                    outy[i,12]<-MBCsamec<-(as.numeric(DNAbias)*(GRmean*totalDNA*FBratio*MBCDNAc+GRmean*totalDNA*(1-FBratio)*MBCDNAc))/TrueMBCsame
                    outy[i,13]<-MBCsamedc<-(as.numeric(DNAbias)*(GRmean*totalDNA*FBratio*as.numeric(DNAexteffF)*MBCDNAdc+GRmean*totalDNA*(1-FBratio)*as.numeric(DNAexteffB)*MBCDNAdc))/TrueMBCsame
                    
                    #True MBC growth if bacteria grow at 0.12 day-1 and fungi at 0.0088 day -1
                    
                    TrueMBCdiff<-(as.numeric(GRfun)*totalDNA*FBratio*as.numeric(MBCDNAratF)+as.numeric(GRbact)*totalDNA*(1-FBratio)*as.numeric(MBCDNAratB))
                    #observed MBC growth if bacteria grow at 0.12 day-1 and fungi at 0.0088 day -1, under various extraction biases
                    outy[i,14]<-MBCdiffd<-(as.numeric(DNAbias)*(as.numeric(GRfun)*totalDNA*FBratio*as.numeric(DNAexteffF)*MBCDNAd+as.numeric(GRbact)*totalDNA*(1-FBratio)*as.numeric(DNAexteffB)*MBCDNAd))/TrueMBCdiff
                    outy[i,15]<-MBCdiffc<-(as.numeric(DNAbias)*(as.numeric(GRfun)*totalDNA*FBratio*MBCDNAc+as.numeric(GRbact)*totalDNA*(1-FBratio)*MBCDNAc))/TrueMBCdiff
                    outy[i,16]<-MBCdiffdc<-(as.numeric(DNAbias)*(as.numeric(GRfun)*totalDNA*FBratio*as.numeric(DNAexteffF)*MBCDNAdc+as.numeric(GRbact)*totalDNA*(1-FBratio)*as.numeric(DNAexteffB)*MBCDNAdc))/TrueMBCdiff
                    
                    outy[i,1]<-MBCDNAratB
                    outy[i,2]<-MBCDNAratF
                    outy[i,3]<-DNAexteffF
                    outy[i,4]<-DNAexteffB
                    outy[i,5]<-MBCexteffF
                    outy[i,6]<-MBCexteffB
                    outy[i,7]<-DNAbias
                    outy[i,8]<-GRbact
                    outy[i,9]<-GRfun
                    outy[i,10]<-FBratio
                    i<-i+1
                  } }}}}}}}}}


outy<-as.data.frame(outy)
outy<-subset(outy, !is.na(outy$V1))
names(outy)<-c("MBCDNAratB", "MBCDNAratF", "DNAexteffF","DNAexteffB","MBCexteffF","MBCexteffB","DNAbias", "GRbact", "GRfun", "FBratio", "MBCsamed", "MBCsamec", "MBCsamedc",  "MBCdiffd","MBCdiffc","MBCdiffdc")

outy0.2FB<-subset(outy, FBratio==0.2)
outy0.3FB<-subset(outy, FBratio==0.3)
outy0.4FB<-subset(outy, FBratio==0.4)
outy0.5FB<-subset(outy, FBratio==0.5)
outy0.6FB<-subset(outy, FBratio==0.6)
outy0.7FB<-subset(outy, FBratio==0.7)
outy0.8FB<-subset(outy, FBratio==0.8)
outy0.9FB<-subset(outy, FBratio==0.9)
outy0.01FB<-subset(outy, FBratio==0.01)
outy0.05FB<-subset(outy, FBratio==0.05)
outy0.95FB<-subset(outy, FBratio==0.95)
outy0.99FB<-subset(outy, FBratio==0.99)
outy0.25FB<-subset(outy, FBratio==0.25)


reference2 <- c( 13.22,1070,0.33,0.5, 0.33,0.32,0.7,0.167,0.0088)#defining parameters to keep for all runs
outysubs0.2FB<- outy0.2FB[apply(outy0.2FB[,1:9], 1, function(x) sum(reference2  == x) >= 8), ]   #subsetting so only get rows where a single value is changed           
outysubs0.3FB<- outy0.3FB[apply(outy0.3FB[,1:9], 1, function(x) sum(reference2  == x) >= 8), ]   #subsetting so only get rows where a single value is changed           
outysubs0.4FB<- outy0.4FB[apply(outy0.4FB[,1:9], 1, function(x) sum(reference2  == x) >= 8), ]   #subsetting so only get rows where a single value is changed           
outysubs0.5FB<- outy0.5FB[apply(outy0.5FB[,1:9], 1, function(x) sum(reference2  == x) >= 8), ]   #subsetting so only get rows where a single value is changed           
outysubs0.6FB<- outy0.6FB[apply(outy0.6FB[,1:9], 1, function(x) sum(reference2  == x) >= 8), ]   #subsetting so only get rows where a single value is changed           
outysubs0.7FB<- outy0.7FB[apply(outy0.7FB[,1:9], 1, function(x) sum(reference2  == x) >= 8), ]   #subsetting so only get rows where a single value is changed           
outysubs0.8FB<- outy0.8FB[apply(outy0.8FB[,1:9], 1, function(x) sum(reference2  == x) >= 8), ]   #subsetting so only get rows where a single value is changed           
outysubs0.9FB<- outy0.9FB[apply(outy0.9FB[,1:9], 1, function(x) sum(reference2  == x) >= 8), ]   #subsetting so only get rows where a single value is changed           
outysubs0.01FB<- outy0.01FB[apply(outy0.01FB[,1:9], 1, function(x) sum(reference2  == x) >= 8), ]   #subsetting so only get rows where a single value is changed           
outysubs0.05FB<- outy0.05FB[apply(outy0.05FB[,1:9], 1, function(x) sum(reference2  == x) >= 8), ]   #subsetting so only get rows where a single value is changed           
outysubs0.25FB<- outy0.25FB[apply(outy0.25FB[,1:9], 1, function(x) sum(reference2  == x) >= 8), ]   #subsetting so only get rows where a single value is changed           
outysubs0.95FB<- outy0.95FB[apply(outy0.95FB[,1:9], 1, function(x) sum(reference2  == x) >= 8), ]   #subsetting so only get rows where a single value is changed           
outysubs0.99FB<- outy0.99FB[apply(outy0.99FB[,1:9], 1, function(x) sum(reference2  == x) >= 8), ]   #subsetting so only get rows where a single value is changed           

outylog0.2FB<-log10(outysubs0.2FB)#taking logarithm of all the values in the table
outylog0.2FB2<-unique( outylog0.2FB )
outylog0.3FB<-log10(outysubs0.3FB)#taking logarithm of all the values in the table
outylog0.3FB2<-unique( outylog0.3FB )
outylog0.4FB<-log10(outysubs0.4FB)#taking logarithm of all the values in the table
outylog0.4FB2<-unique( outylog0.4FB )
outylog0.5FB<-log10(outysubs0.5FB)#taking logarithm of all the values in the table
outylog0.5FB2<-unique( outylog0.5FB )
outylog0.6FB<-log10(outysubs0.6FB)#taking logarithm of all the values in the table
outylog0.6FB2<-unique( outylog0.6FB )
outylog0.7FB<-log10(outysubs0.7FB)#taking logarithm of all the values in the table
outylog0.7FB2<-unique( outylog0.7FB )
outylog0.8FB<-log10(outysubs0.8FB)#taking logarithm of all the values in the table
outylog0.8FB2<-unique( outylog0.8FB )
outylog0.9FB<-log10(outysubs0.9FB)#taking logarithm of all the values in the table
outylog0.9FB2<-unique( outylog0.9FB )
outylog0.01FB<-log10(outysubs0.01FB)#taking logarithm of all the values in the table
outylog0.01FB2<-unique( outylog0.01FB )
outylog0.05FB<-log10(outysubs0.05FB)#taking logarithm of all the values in the table
outylog0.05FB2<-unique( outylog0.05FB )
outylog0.25FB<-log10(outysubs0.25FB)#taking logarithm of all the values in the table
outylog0.25FB2<-unique( outylog0.25FB )
outylog0.95FB<-log10(outysubs0.95FB)#taking logarithm of all the values in the table
outylog0.95FB2<-unique( outylog0.95FB )
outylog0.99FB<-log10(outysubs0.99FB)#taking logarithm of all the values in the table
outylog0.99FB2<-unique( outylog0.99FB )


write.table(outylog0.2FB2, "~/Desktop/LuizMicrocosms/CUEsensitivityAnalysis190207log100.2FB.txt", sep="\t",row.names = FALSE)
write.table(outylog0.3FB2, "~/Desktop/LuizMicrocosms/CUEsensitivityAnalysis190207log100.3FB.txt", sep="\t",row.names = FALSE)
write.table(outylog0.4FB2, "~/Desktop/LuizMicrocosms/CUEsensitivityAnalysis190207log100.4FB.txt", sep="\t",row.names = FALSE)
write.table(outylog0.5FB2, "~/Desktop/LuizMicrocosms/CUEsensitivityAnalysis190207log100.5FB.txt", sep="\t",row.names = FALSE)
write.table(outylog0.6FB2, "~/Desktop/LuizMicrocosms/CUEsensitivityAnalysis190207log100.6FB.txt", sep="\t",row.names = FALSE)
write.table(outylog0.7FB2, "~/Desktop/LuizMicrocosms/CUEsensitivityAnalysis190207log100.7FB.txt", sep="\t",row.names = FALSE)
write.table(outylog0.8FB2, "~/Desktop/LuizMicrocosms/CUEsensitivityAnalysis190207log100.8FB.txt", sep="\t",row.names = FALSE)
write.table(outylog0.9FB2, "~/Desktop/LuizMicrocosms/CUEsensitivityAnalysis190207log100.9FB.txt", sep="\t",row.names = FALSE)
write.table(outylog0.01FB2, "~/Desktop/LuizMicrocosms/CUEsensitivityAnalysis190207log100.01FB.txt", sep="\t",row.names = FALSE)
write.table(outylog0.05FB2, "~/Desktop/LuizMicrocosms/CUEsensitivityAnalysis190207log100.05FB.txt", sep="\t",row.names = FALSE)
write.table(outylog0.25FB2, "~/Desktop/LuizMicrocosms/CUEsensitivityAnalysis190207log100.25FB.txt", sep="\t",row.names = FALSE)
write.table(outylog0.95FB2, "~/Desktop/LuizMicrocosms/CUEsensitivityAnalysis190207log100.95FB.txt", sep="\t",row.names = FALSE)
write.table(outylog0.99FB2, "~/Desktop/LuizMicrocosms/CUEsensitivityAnalysis190207log100.99FB.txt", sep="\t",row.names = FALSE)
